# Targeting CD301+ macrophage inhibits endometrial fibrosis and improves pregnancy outcome

**DOI:** 10.1101/2023.06.09.544336

**Authors:** Haining Lv, Haixiang Sun, Limin Wang, Simin Yao, Dan Liu, Xiwen Zhang, Zhongrui Pei, Jianjun Zhou, Huiyan Wang, Jianwu Dai, Guijun Yan, Lijun Ding, Zhiyin Wang, Chenrui Cao, Guangfeng Zhao, Yali Hu

## Abstract

Macrophages are a key and heterogeneous cell population involved in endometrial repair and regeneration during the menstrual cycle, but their role in the development of intrauterine adhesion (IUA) and sequential endometrial fibrosis remains unclear. Here, we reported that CD301+ macrophages were significantly increased and showed their most active interaction with profibrotic cells in the endometria of IUA patients compared with the normal endometria by single-cell RNA sequencing, bulk RNA sequencing and experimental verification. Increasing CD301+ macrophages promoted the differentiation of endometrial stromal cells into myofibroblasts and resulted in extracellular matrix accumulation, which destroyed the physiological architecture of endometrial tissue, drove endometrial fibrosis and ultimately led to female infertility or adverse pregnancy outcomes. Mechanistically, CD301+ macrophages secreted GAS6 to activate the AXL/NF-κB pathway, up-regulating the profibrotic protein synthesis. Targeted deletion of CD301+ macrophages or inhibition of AXL by Bemcentinib blunted the pathology and improved the outcomes of pregnancy in mice, supporting the therapeutic potential of targeting CD301+ macrophages for treating endometrial fibrosis.

## Introduction

Intrauterine adhesion (IUA), characterized by endometrial fibrosis, is a common cause of the refractory uterine infertility, which is a clinically devastating condition for the patients who desire to have children. About ninety percent of IUA is secondary to uterine injury, and a few are associated with severe endometrial infections, especially tuberculous endometritis (de Miguel-Gómez, et al.,2021; Freedman and Schlaff,2021; Song, et al.,2021). Currently routine hysteroscopic adhesiolysis shows high recurrence of IUA (Hanstede, et al.,2015) and may not ameliorate the endometrial fibrosis. Hence, seeking a therapeutic target for IUAs and endometrial fibrosis is urgently needed. Myofibroblasts, characterized by increased expression of collagen 1 and α-SMA, are important effector cells for IUAs, which produce a large amount of extracellular matrix (ECM) in the endometria (Rodriguez, et al.,2019; Zhu, et al.,2019). Myofibroblasts mainly differentiate from stromal cells or fibroblasts upon stimulation by various cytokines in the fibrotic microenvironment (Liu, et al.,2021; Vorstandlechner, et al.,2021; Yang, et al.,2021).

It has been reported that the pathological process of tissue fibrosis in the lung, kidney and liver is intimately related to injury and a persistent inflammatory response by activation of immune cells. These immune cells release cytokines and other inflammatory mediators to eliminate the initial insults, produce appropriate ECM, repair the lesions and finally restore local homeostasis. However abnormal activation of detrimental subtypes of immune cells or failure to inhibit the production of profibrotic factors will exacerbate the deposition of ECM and promote aberrant scar development (Ramachandran, et al.,2019; Tang and Yiu,2020; Liu, et al.,2021). Previous study showed that during menstrual period, endometrial macrophage plays an important role in removing endometrial debris and producing a variety of inflammatory and growth factors to promote the repair, regeneration and growth of endometrial tissue (Garry, et al.,2010;

Thiruchelvam, et al.,2013). In contrast, their dysregulation is linked to the pathogenesis of some endometrial diseases, such as endometriosis (Hogg, et al.,2021) and endometrial cancer (Ning, et al.,2016), while their role in the fibrosis of IUA has been rarely reported. In other fibrotic diseases, the function of macrophages always acts as a double-edged sword depending on their subtypes and the local tissue microenvironment, therefore many studies are exploring to determine the key subtypes of macrophages that determine the fate of fibrosis in the different organs (Shook, et al.,2018; Chakarov, et al.,2019; Zindel, et al.,2021).

Here, we reported that CD301+macrophages were the most active subtypes of macrophages to communicate with the profibrotic cells by 10x Genomics single-cell RNA sequencing (scRNA-seq) on endometria. We further revealed that CD301+ macrophages were significantly increased and promoted stromal cells to differentiate into myofibroblasts through the GAS6/AXL/NF-κB signaling pathway of IUA. Deleting CD301+ macrophages or inhibiting the activation of AXL using Bemcentinib reversed endometrial fibrosis both *in vitro* and *in vivo*, and most importantly, it can significantly improve the pregnancy outcome. Overall, our research deciphered the function and mechanism of CD301+ macrophages in endometrial fibrosis of IUA and provided potential targets for developing novel therapeutic interventions for this disease.

## Results

### CD301+ macrophages show the most active interaction with profibrotic myofibroblast-like cells in IUA

To investigate the cellular features of endometrial cells in fibrotic endometria of IUA patients at single-cell resolution, scRNA-seq by 10x Genomics was applied in three IUA endometria and normal endometria respectively (Appendix Table S1). After stringent cell filtration and batch correction, graph-based clustering was first performed for IUA and normal endometria to catalog a total of 57,770 cells into nine distinct clusters identified by exploration of cluster marker genes that were compared to annotated major cell types in reported endometrial atlases (Wang, et al.,2020; Lv, et al.,2022), including stromal cells, epithelial cells, endothelial cells, myofibroblast-like cells, natural killer cells, T cells, mast cells, B cells and macrophages/dendritic cells (DCs) in normal and IUA endometria (Fig 1A, Fig S1A, Appendix Table S2). The UMAP distributions of markers for each type of cells were shown in Fig S1B. Then we captured the macrophage/ DC and used higher resolution to identify them as two types of cells (macrophage, DC) (Fig 1B, Fig S1C, Appendix Table S3). We used CD14, CD68 and CD163 to identify endometrial macrophage and CD1C, FCER1A and CD1E to identify endometrial DC (Fig 1C). Moreover, several subtypes of stromal cell, epithelial cell and endothelial cell were also identified, and their markers and the percentage of each cluster between controls and patients were shown in Appendix Fig. S2A-I.

**Figure 1.**
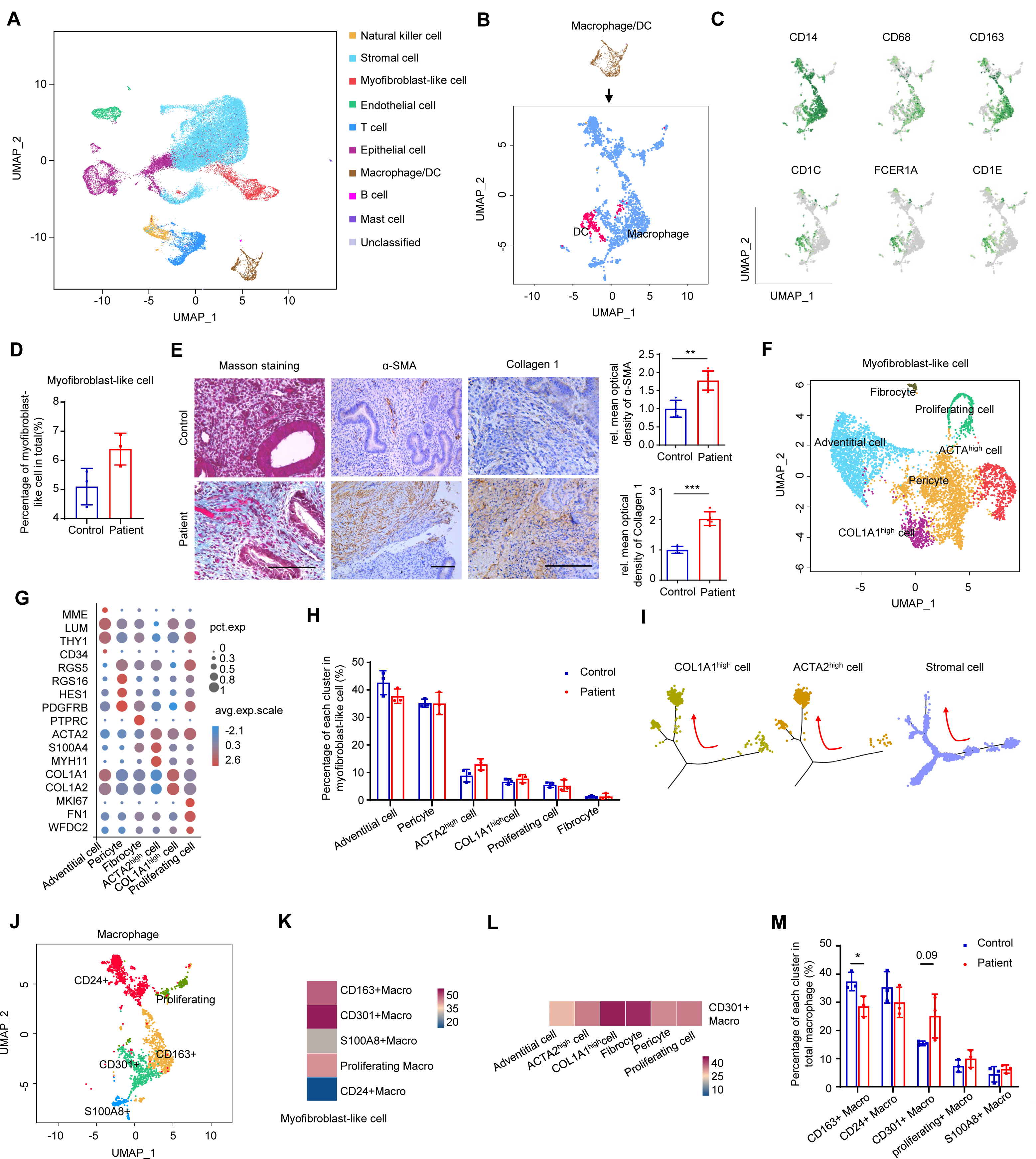
scRNA-seq cartography of myofibroblast-like cell and macrophage in human endometrium between IUA patients and healthy controls. A. The UMAP plot displaying total cells from controls and patients with IUA (n=6 samples). B. The UMAP plot displaying dendritic cells (DC) and macrophages (n=6 samples). C. The UMAP plot showing the markers for macrophages and DC (n=6 samples). D. The percentage of myofibroblast-like cells in total endometrial cells between controls and patients with IUA (n=3 samples for each group). E. Masson trichrome staining and immunostaining of α-SMA and Collagen 1 in endometrium of controls and patients with IUA (n=5 samples for each group). Scale bars: 100μm. F. The UMAP plot embedding subpopulations of myofibroblast-like cells annotated by cell type/state from controls and patients with IUA (n=6 samples). G. Expression of marker genes for each cell subtype of myofibroblast-like cells (n=6 samples). H. The percentage of each cell subtype in total myofibroblast-like cells between controls and patients with IUA (n=3 samples for each group). I. The branched trajectory of state transition of COL1A1^high^ cells, ACTA2^high^ cells and stromal cells in a two-dimensional state space inferred by Monocle (n=6 samples). J. The UMAP plot showing subtypes of macrophages from controls and patients with IUA (n=6 samples). K. The abundance of connection between subtypes of macrophages (as legends’ cell) and myofibroblast-like cells (as receptors’ cell) utilizing CellPhoneDB (n=6 samples). L. The abundance of connection between CD301+ macrophages (as legends’ cells) and subtypes of myofibroblast-like cells (as receptors’ cells) utilizing CellPhoneDB (n=6 samples). M. The percentage of each cell subtype of macrophages between controls and patients with IUA (n=3 samples for each group). Macro, macrophage. Data are presented as mean ±SEM. (D, F, H, and M) Two-tailed Student’s t-test. *, P<0.05; **, P<0.01; ***, P<0.001.

We next focused on the fibrotic changes in the endometria of IUA patients. The proportion of myofibroblast-like cell was increased (Fig 1D) and the upregulation of α-SMA and Collagen 1 expression as well as Masson trichrome staining showed much more collagen deposition in the endometrial stroma, compared to normal endometria (Fig 1E), suggesting significant fibrotic changes in endometria of IUA. As myofibroblast plays a pivotal role in tissue fibrosis (Valenzi, et al.,2019), we captured this type of cells and subclustered them into adventitial cell, pericyte, fibrocyte, ACTA2^high^ cell, proliferating cell and COL1A^high^ cell according to the most upregulated genes and previously reported marker genes (Fig 1F-G, Appendix Fig S1D) (Dupuis and Kern,2014; Dong, et al.,2015; Falke, et al.,2015; Pilling, et al.,2015; Kirkwood, et al.,2021; Zhu, et al.,2021). Marker genes, LUM in the adventitial cell and PDGFRB in the pericyte were verified in Appendix Fig S1E. The proportions of ACTA2^high^ and COL1A1^high^ cell in IUA endometria were higher than those in normal endometria, verified by immunostaining of α-SMA and Collagen 1 (Fig 1E and H). Pseuduotime analysis of stromal cell and myofibroblast-like cell inferred that profibrotic myofibroblast-like cell---COL1A1^high^ cell and ACTA2^high^ cell were derived from stromal cell (Fig 1I).

To explore the effect of endometrial macrophage on IUA pathogenesis, we captured macrophage and subclustered them into five populations, CD163+ macrophage, CD301+ macrophage (whose encoding gene is CLEC10A), CD24+ macrophage, S100A8+ macrophage and proliferating macrophage which were named according to their specific highly expressed genes and previously reported marker genes (Jensen, et al.,2012; Shook, et al.,2018; Ganta, et al.,2019) (Fig 1J, Appendix Fig S3A). Marker genes, CD163, CD24 and CD301 were verified by immunostaining in Appendix Fig S1E and Fig 2B. CD163+ endometrial macrophage played an important role in host defense and the regulation of tissue homeostasis including tissue breakdown, clearance, and angiogenic remodeling (Jensen, et al.,2012). CD24 interacted with Siglec-10 on innate immune cells to dampen damaging inflammatory responses to infection (Barkal, et al.,2019). CD24+ macrophage also expressed high-level of GATA6 shown in Appendix Fig S3A which was reported to have a phenotype for reparative immune response and anti-fibrotic functions (Deniset, et al.,2019). CD301+ macrophage was reported to promote skin wound repair (Shook, et al.,2018) but be a key driver of autoimmune cardiac valve inflammation and fibrosis (Meier, et al.,2018) while S100A8+ macrophage was previously reported to inhibit endothelial angiogenic potential in a paracrine manner (Ganta, et al.,2019). To confirm which subtype of macrophage had a preponderant effect on the myofibroblast-like cells in the endometria from IUA patients, we used CellPhoneDB analysis, a public repository of ligands, receptors and their interactions, to enable a comprehensive, systematic analysis of cell-cell communication molecules to analyze the connections between subtypes of macrophages and myofibroblast-like cells. The abundance of connections between each subtype of macrophage as senders and myofibroblast-like cells as receivers showed that CD301+macrophage sent much more signaling to myofibroblast-like cells than others in normal and IUA endometria (Fig 1K). Moreover, we explored the relationship between subtypes of myofibroblast-like cells and CD301+ macrophages and the abundance of connections indicated that the most active receivers of CD301+ macrophage were COL1A1^high^ cell, fibrocyte and ACTA2^high^ cell in the endometrium (Fig 1L). It has previously been shown that fibrocyte can be derived from monocyte differentiation into fibroblast-like cells and contribute to fibrotic diseases(Pilling, et al.,2015), and COL1A1^high^ cell and ACTA2^high^ cell secretes much more extracellular matrix and fibrotic cytokines to further activate fibroblasts to differentiate into myofibroblasts in tissues, which is the central link to fibrosis. The close connection between CD301+macrophage and pro-fibrotic cells in our analysis suggested that CD301+macrophage might be the key subpopulation of macrophage influencing the differentiation and survival of myofibroblasts. The up-regulated expression of fibrosis-related genes, such as TGFB1, TIMP1 and NFKB1 and inflammation-related genes, CXCL8, IL1B and IL18 in CD301+ macrophage verified by qRT-PCR, further indicated that CD301+ macrophage had the deleterious role in IUA (Appendix Fig S3B-C). The signaling of upregulated genes which was enriched in CD301+ macrophages of the patients compared to controls by KEGG analysis suggested that CD301+ macrophages participated in more inflammatory activities in patients (Appendix Fig S3D). CellPhone DB analysis among other types of cells, such as epithelial cells, stromal cells, macrophages, and their progenitor-like cells were shown in Appendix Fig. S2J-L.

**Figure 2.**
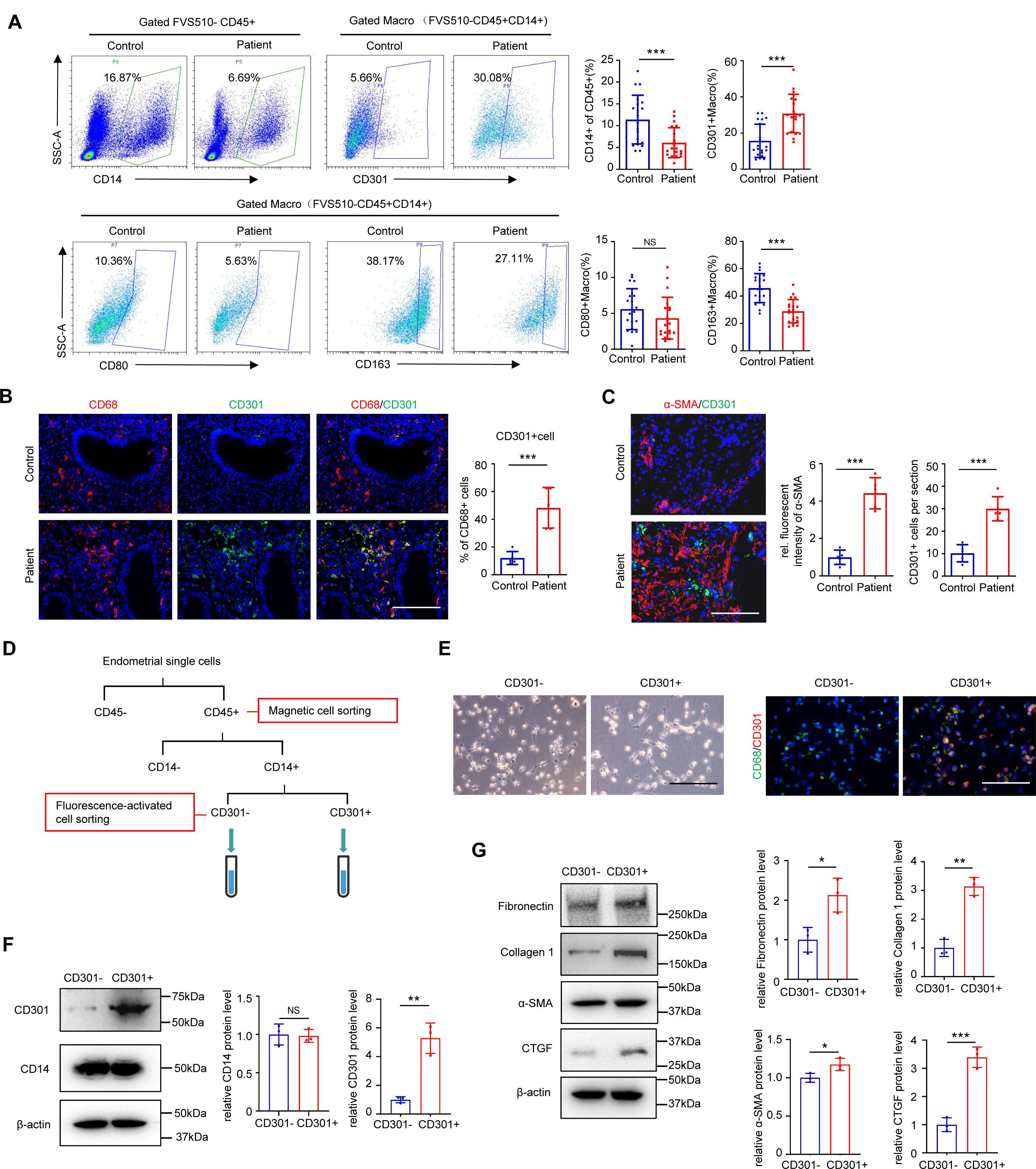
The changes of endometrial CD301+ macrophage in IUA and its effect on primary human endometrial stromal cells. A. Flow cytometric analysis of the proportion of CD14+ cells, CD301+ macrophages, CD80+ macrophages and CD163+ macrophages in the endometrium of normal controls and patients with IUA. Macro, macrophage (n=20 samples for each group). B. Representative immunostaining images of CD301 and CD68 from samples of normal controls and patients with IUA (n=5 samples for each group). C. Localization of CD301 and α-SMA in endometrium of IUA by immunofluorescence (n=5 samples for each group). D. Schema describing the approach to sort CD301-macrophages and CD301+ macrophages from human endometrial single cells. E. Representative images from CD301-macrophages and CD301+ macrophages. Left panel displays the morphology of CD301-macrophages and CD301+ macrophages and right panel shows the expression of CD68 and CD301 by immunofluorescence. F. The protein level of CD14 and CD301 in CD301-macrophages and CD301+ macrophages examined by western blot (n=3 technical replicates for each group). G. The protein level of Fibronectin, Collagen1, α-SMA and CTGF expression in primary human endometrial stromal cells (hESCs) treated by the supernatants of CD301- and CD301+ macrophages for 48h, respectively examined by western blot (n=3 technical replicates for each group). Scale bar: 100μm. Data are presented as mean ±SEM. (A, B, C, F, and G) Two-tailed Student’s t-test. *, P<0.05; **, P<0.01; ***, P<0.001. NS, not significant.

Furthermore, we applied CellChat analysis to investigate cellular cross-talk between CD301+ macrophage and pro-fibrotic myofibroblast-like cells in both controls and patients. The results indicated that CD301+ macrophage was more active in TWEAK signaling pathway which targeted on ACTA2^high^ cell and COL1A1^high^ cell, and IGF signaling pathway which specifically targeted on fibrocyte, especially in patients (Appendix Fig S3E-H). Both TWEAK and IGF signaling pathway are recognized to positively regulate cell proliferation and growth (Fang, et al.,2022; Lv, et al.,2022), suggesting that CD301+ macrophage participated in the increase of pro-fibrotic myofibroblast-like cells in patients. Additionally, the change tendency of each macrophage cluster showed that CD301+ macrophages accounted for a higher percentage in IUA endometria than in normal endometria (Fig 1M).

### CD301+ macrophages are significantly increased and promote stromal cells differentiation into myofibroblasts in the endometria of IUA

To verify the functions and changes in CD301+ macrophages in IUA analyzed by scRNA-seq, we used flow cytometry to examine the changes in the proportion of macrophages in the endometria of IUA patients and controls. The results showed that the proportion of CD301+ macrophages was significantly increased along with the decreases in total macrophages (CD14+CD45+ cells) and CD163+ macrophages (Fig 2A, Appendix Fig S4A-B). The lower number of CD14+CD45+ cells might contribute to the insufficient proliferation and growth of endometrial stromal cells which we documented previously (Lv, et al.,2022). Consistently, the decreased number of proliferating stromal cells were also observed in IUA patients shown in Appendix Fig S2C. The immunostaining further showed that CD301 was co-stained with pan-macrophage marker, CD68, and up-regulated in IUA endometria (Fig 2B).

We next examined the relationship between CD301+ macrophages and endometrial fibrosis. Immunofluorescence detection revealed that there were more CD301+ macrophages near fibrotic regions marked by α-SMA, but fewer CD301+ macrophages in α-SMA negative-region (Fig 2C). To explore the role of CD301+ macrophages directly, we digested and isolated human endometrial cells to sort immune cells by CD45 magnetic beads, and then obtained CD301-and CD301+ macrophages by flow sorting (Fig 2D). Representative images of CD301- and CD301+ macrophages were shown in Fig 2E. The purity of sorted cells was identified by co-staining of CD68 and CD301 and western blot examination of CD14 and CD301 protein expression (Fig 2E and F). As profibrotic myofibroblast-like cells---COL1A1^high^ cells and ACTA2^high^ cells were differentiated from stromal cells shown in Fig 1I, we then used primary human endometrial stromal cells (hESCs) *in vitro* to investigate the role of CD301+ macrophages in myofibroblast differentiation. The results showed that the supernatants of CD301+ macrophages significantly enhanced the expression of fibrotic markers, Fibronectin, Collagen1, α-SMA and CTGF in hESCs indicating that there may exist stromal cells-myofibroblasts transition (Fig 2G). No significant difference was found between the group of hESCs treated with CD301-macrophage and the group of hESCs only (Appendix Fig S4C). The results above demonstrated that CD301+ macrophages were significantly increased in IUA and promoted pro-fibrotic differentiation of stromal cells.

### CD301+ macrophages activate the AXL receptor in hESCs by secreting GAS6

To understand the molecular mechanisms by which CD301+ macrophages to promote hESCs differentiation to myofibroblasts, we tested the roles of the cytokines secreted from CD301+ macrophages suggested by the GSE105789 dataset, which analyzed the differential expression of genes between CD301b+ macrophages (the ortholog of CD301+ macrophages in mice) and F4/80-immune cells(Shook, et al.,2018). Gas6, Ccl8. Sepp1, Apoe and Pltp were found to be the most up-regulated genes which encoded secretory proteins in CD301b+ macrophages (Fig 3A). Our qRT-PCR results confirmed that two times higher expression of GAS6 and CCL8 in CD301+ macrophages than in CD301-macrophages from human endometria at the mRNA level (Fig 3B) and the results of GAS6 and CCL8 expression at the protein level further confirmed that GAS6 was significantly increased in CD301+ macrophages of endometrium (Fig 3C). Immunostaining showed that the expression of GAS6 was co-stained with CD301 and was upregulated in the endometria of IUA patients (Fig 3D). Together, these data suggested that GAS6 was an important molecule secreted by CD301+ macrophages to function.

**Figure 3.**
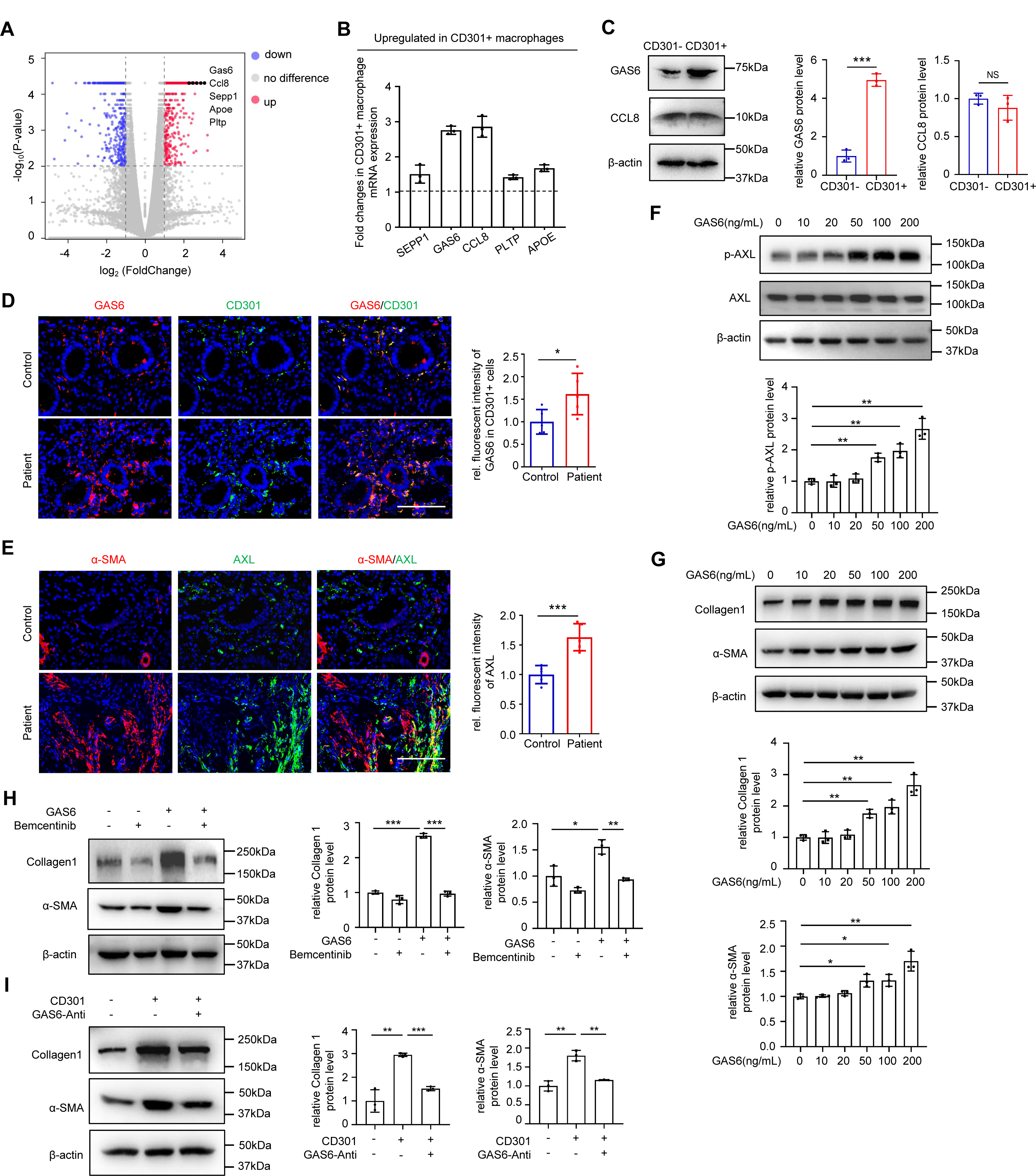
CD301+ macrophage activates GAS6/AXL signaling to promote α-SMA and Collagen1 expression in hESCs. A. Volcano plots of genes between CD301b+ macrophage and F4/80-immune cells based on the reanalysis of a published dataset (GSE105789). The labeled genes which encoded secretory proteins were selected conformed to fragments per kilobase per million (FPKM)>100 in CD301+ macrophage, at least a factor of 1.5 and false discovery rate of <0.0001. B. qRT-PCR of SEPP1, GAS6, CCL8, PLTP and APOE expression in human endometrial CD301+ macrophages compared to CD301-macrophages from magnetic cell sorting combined with fluorescence-activated cell sorting. Black lines indicated expression levels in CD301-macrophage which had been normalized to 1, and the bar level indicated fold upregulated changes of gene expression levels in CD301+ macrophage compared to CD301-macrophage (n=3 biological replicates for each group). C. The protein level of GAS6 and CCL8 expression in sorted human endometrial CD301- and CD301+ macrophage (n=3 technical replicates for each group). D. Localization of GAS6 and CD301 in endometrium of normal controls and patients with IUA examined by immunofluorescence (n=5 samples for each group). E. Localization of AXL and α-SMA in endometrium of normal controls and patients with IUA examined by immunofluorescence (n=5 samples for each group). F. Western blot analysis of phospho-AXL (p-AXL) and AXL in hESCs treated with GAS6 at dose of 10, 20, 50, 100 and 200 ng/mL for 72h (n=3 technical replicates for each group). G. The protein level of Collagen1 and α-SMA expression in hESCs after treatment with GAS6 at dose of 10, 20, 50, 100 and 200 ng/mL for 72h examined by western blot (n=3 technical replicates for each group). H. Western blot analysis of α-SMA and Collagen1 in hESCs treated with Bemcentinib (1μM) for 1 hour, followed by GAS6 (50ng/mL) for 72h (n=3 technical replicates for each group). I. Western blot analysis of α-SMA and Collagen1 in hESCs treated with the supernatants of CD301- and CD301+ macrophages and GAS6 neutralizing antibody (2.5μg/mL) for 72h (n=3 technical replicates for each group). Scale bar: 100μm. Data are presented as mean ±SEM. (C, D, and E) Two-tailed Student’s t-test. (F, G, H, and I) One-way ANOVA with Tukey’s post hoc analysis. *, P<0.05; **, P<0.01; ***, P<0.001. NS, not significant.

The receptor of GAS6 is a member of the TAM family of receptor tyrosine kinases, including TYRO3, AXL and MERTK, and among them, GAS6 has the highest affinity for AXL(Kariolis, et al.,2017). Thus, we first investigated the role of AXL in the endometrium. The expression of AXL obviously increased in the IUA and most of AXL was located in fibrotic regions marked by α-SMA (Fig 3E). To further evaluate the GAS6-AXL link, we studied the effect of GAS6 on AXL activation in hESCs *in vitro*. The mRNA level of AXL was upregulated after treatment with GAS6 for 72 h (Appendix Fig S5A), and western blot analysis showed that GAS6 treatment led to an increase in the protein levels of phospho-AXL in a dose-and time-dependent manner (Fig 3F and Appendix Fig S5B). To explore whether the two other receptors MERTK and TYRO3 could also be activated by GAS6 treatment in the endometrium, we examined their expression and location in normal and IUA endometria and their activated level in hESCs after treatment with GAS6. The results showed that the expression levels of MERTK and TYRO3 were not significantly different in endometria between IUA patients and controls (Appendix Fig S5C) and there were no obvious changes in phospho-MERTK, MERTK, phospho-TYRO3 and TYRO3 expression in hESCs under GAS6 stimulation (Appendix Fig S5D-E). Taken together, these data demonstrated that CD301+ macrophages could produce GAS6 to specifically activate AXL in hESCs.

### The activation of AXL induces the up-regulation of profibrotic genes in endometrial stromal cells

To investigate the role of AXL in endometria of IUA, we used GAS6 to treat hESCs to activate AXL and found that Collagen l and α-SMA at both the mRNA and protein levels were significantly increased in hESCs (Fig 3G, Appendix Fig S5F-H). In addition, GAS6 treatment led to the increased expression of Collagen l and α-SMA in hESCs in a dose- and time-dependent manner (Fig 3G, Appendix Fig S5G-H). Then, we inhibited AXL with Bemcentinib (MedChemExpress) in the supernatants of hESCs treated with GAS6. During dose exploration of Bemcentinib, we found that 1μM was a better dose to examine AXL activation and downstream Collagen1 expression (Appendix Fig S6A), so we used 1μM to do the following *in vitro* experiments. It was shown that the expression levels of Collagen 1 and α-SMA were significantly reduced after stimulation with Bemcentinib (Fig3H, Appendix Fig S6B), indicating that AXL pathway was important for GAS6 functioning. We next added a GAS6 antibody for neutralization in the supernatants of CD301+ macrophages and put their supernatants into hESCs. The results showed that supernatants from CD301+ macrophages plus GAS6 antibody could obviously inhibit the expression of Collagen 1 and α-SMA in hESCs at both the mRNA and protein levels compared to the treatment with supernatants only from CD301+ macrophages (Fig 3I, Appendix Fig S6C-D). These findings demonstrated that GAS6/AXL signaling had a pivotal role in promoting stromal cells to differentiate into myofibroblasts in the endometrium.

### NF-κB signaling activation is essential for myofibroblast differentiation induced by GAS6/AXL

To further decipher the molecular mechanisms and signaling pathways in GAS6/AXL-induced myofibroblast differentiation, we compared the transcriptomes of hESCs treated with GAS6 or without GAS6 by bulk RNA sequencing. The KEGG analysis showed that the significantly upregulated genes in hESCs treated with GAS6 were most enriched in NF-κB signaling, which has been reported to promote fibrogenesis in many organs(Oakley, et al.,2009; Moles, et al.,2013; Chen, et al.,2017) (Fig 4A). RelA/p65, a crucial member of the canonical NF-κB pathway(Lawrence,2009), was validated to have increased phosphorylation in hESCs after treatment with GAS6 (Fig 4B). Furthermore, a GAS6 antibody for neutralization and Bemcentinib for suppressing AXL activation significantly inhibited the phospho-p65 (Fig 4C) and phospho-p65 expression was also upregulated in the endometria of IUA patients *in vivo* (Fig 4D). These results indicated that NF-κB pathway was an important downstream pathway for GAS6/AXL. To further determine its key role in GAS6/AXL-induced myofibroblast differentiation, we used an inhibitor of the NF-κB pathway, JSH23 (MedChemExpress) and found that the expression levels of Collagen 1 and α-SMA were significantly decreased upon treatment with JSH23, followed by GAS6 (Fig 4E-F). Together, these data showed that GAS6/AXL accelerated myofibroblast differentiation by activating the NF-κB signaling pathway in the endometria of IUA.

**Figure 4.**
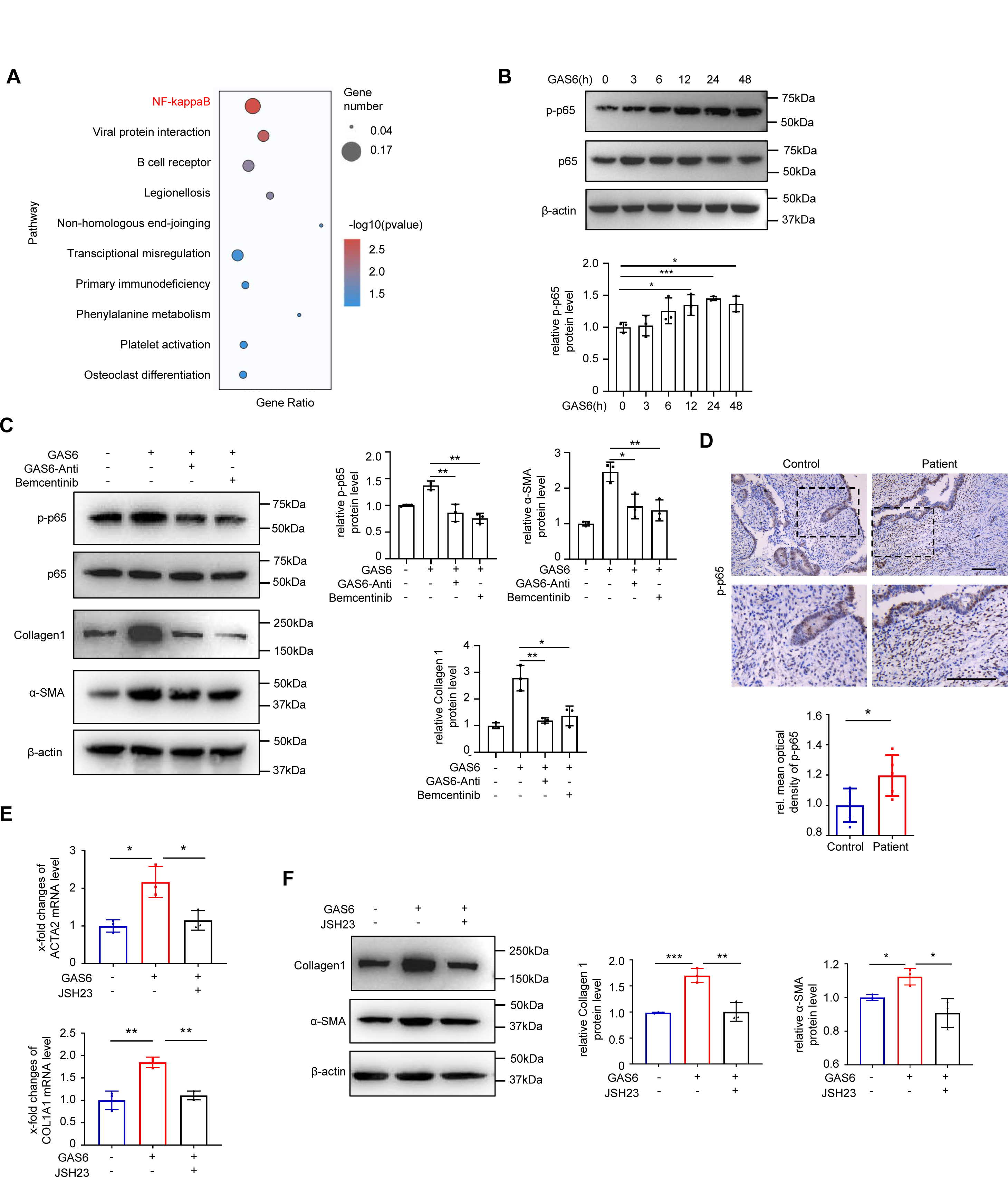
NF-κB signaling pathway is an essential downstream for GAS6/AXL functioning. A. Scatterplot of enriched KEGG pathways statistics was made. Gene ratio is the ratio of the differentially expressed gene number to the total gene number in a certain pathway. The color and size of the dots represent the range of the p-value and the number of differentially expressed genes mapped to the indicated pathways, respectively. Top 10 enriched pathways are shown in the figure. KEGG analysis of bulk RNA-sequencing showing the enrichment of differentially expressed genes between hESCs treated with and without GAS6 (50ng/mL) for 72h (n=3 biological replicates for each group). B. Western blot analysis of phospho-p65 (p-p65) and p65 expression in hESCs treated with GAS6 (50ng/mL) at time of 3, 6, 12, 24 and 48h (n=3 technical replicates for each group). C. Western blot analysis of p-p65, p65, Collagen1 and α-SMA expression in hESCs treated with or without GAS6 neutralizing antibody (2.5μg/mL) or Bemcentinib (1μM) for 1 hour, followed by GAS6 protein (50ng/mL) for 72h (n=3 technical replicates for each group). D. Representative immunostaining images of p-p65 in endometrium of normal controls and patients with IUA (n=5 samples for each group). E. qRT-PCR analysis of ACTA2 and COL1A1 expression in hESCs treated with GAS6 (50ng/mL) or JSH23 (10μM) for 72h (n=3 biological replicates for each group). F. Western blot analysis of Collagen1 and α-SMA expression in hESCs treated with GAS6 (50ng/mL) or JSH23 (10μM) for 72h (n=3 technical replicates for each group). Scale bar: 100μm. Data are presented as mean ±SEM. (B, C, E and F) One-way ANOVA with Tukey’s post hoc analysis. (D) Two-tailed Student’s t-test. *, P<0.05; **, P<0.01; ***, P<0.001.

### The IUA mouse model has similar changes in endometrial CD301+ macrophages to IUA patients

To better understand the functional effects of CD301+ macrophages *in vivo*, we first tried to simulate the process of IUA in murine endometria by curettage. The murine ortholog of CD301 is macrophage galactose-type C-type lectin 2 (*Mgl2*, encoding the CD301b protein). A diagram of this operation in mice was shown in Fig 5A. Masson trichrome staining showed increased collagen deposition, and immunofluorescence staining showed up-regulated Collagen 1 expression in the IUA group (Fig 5B). qRT-PCR further confirmed that the profibrotic myofibroblast markers Acta2 and Col1A1 were significantly increased in the uterus of the IUA group (Fig 5C). The proportion of total macrophages marked by F4/80+CD11b+ was significantly reduced but CD301b+ macrophages were augmented in the uterus of the IUA model tested by flow cytometry. Additionally, CD86 which was mostly used for labeling M1-like macrophage in mouse was increased and CD163, usually labeling M2-like macrophage was decreased. These results have the consistent tendency with those in human. (Fig 5D, Appendix Fig S7A). Furthermore, the expression of GAS6 co-stained with CD301b and AXL was also increased in the IUA models (Fig 5E-F). Overall, these changes in the mouse models were similar to those in the endometria of IUA patients.

**Figure 5.**
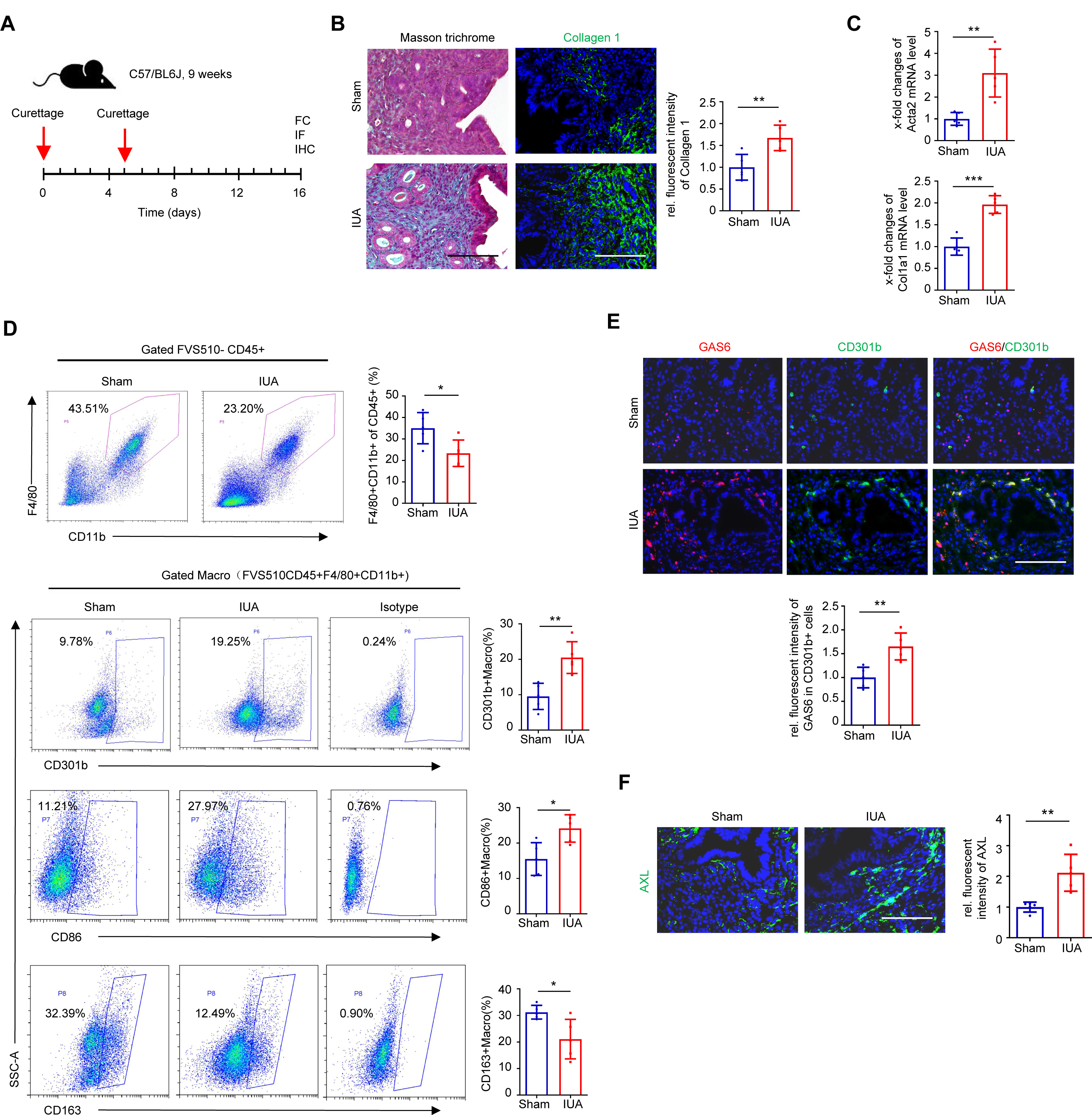
The proportion of CD301b+ macrophage is increased along with the up-regulated GAS6, AXL and Collagen1 expression in IUA model of mice. A. Schema describing the strategy to build the IUA model of mice. B. Representative images of Masson trichrome staining and immunostaining for Collagen 1 between endometrium of Sham and IUA mice (n=5 mice for each group). C. qRT-PCR analysis of Acta2 and Col1a1 expression in uterus of Sham and IUA mice (n=5 mice for each group). D. Flow cytometric analysis of the proportions of CD11b+F4/80+ cells (pan-macrophages) and CD301b+ macrophages in the uterus of Sham and IUA mice (n=5 mice for each group). Macro, macrophage. E. Representative images of immunostaining for GAS6 (red) and CD301b (green) in the endometrium of Sham and IUA mice (n=5 mice for each group). F. Representative images of immunostaining for AXL in the endometrium of Sham and IUA mice (n=5 mice for each group). Scale bar: 100μm. Data are presented as mean ±SEM. (B, C, D and E) Two-tailed Student’s t-test. (F) Mann-Whitney U test. *, P<0.05; **, P<0.01; ***, P<0.001.

### Depletion of CD301b+ macrophages reverses the endometrial fibrosis and restores the pregnancy capability of the IUA model in mice

To further confirm the role of CD301b+ macrophages in the development of endometrial fibrosis *in vivo,* we used Mgl2^+/DTR-GFP^ (Mgl2-DTR) mice, in which CD301b+ macrophages can be specifically and inducibly depleted by injecting diphtheria toxin (DT, *i.p.*)(Kumamoto, et al.,2013). The GFP-positive cells in murine endometrium were confirmed to be mostly depleted for 2, 4 and 6 days after a single DT injection and there was no obvious difference in Masson trichrome staining by intraperitoneal injection of DT four days later (Appendix Fig S7B-C). In addition, the numbers of live pups and the absorbed embryos were also similar between mice with and without DT injection (Appendix Fig S7D) indicating that the simple injection of DT had no impact on the development of endometrial fibrosis or the outcome of pregnancy.

We next performed uterine curettage on Mgl2-DTR mice and injected DT intraperitoneally at 5- and 8-days post operation, respectively. The diagram for this operation on mice was shown in Fig 6A. Flow cytometric analysis showed that both the proportion and the number of CD301b+ macrophages in the uterus were significantly diminished in the DT group (Fig 6B). In line with the results above, the expression of GAS6 in GFP-labeled CD301b+ macrophages in murine endometrium was increased in the IUA model of mice, but they were significantly dampened in the IUA+DT group (Fig 6C). The mRNA levels of Acta2 and Col1a1 were also decreased in the IUA+DT group (Fig 6D). Moreover, the expression levels of AXL, Collagen1 and phospho-p65 were obviously inhibited, accompanied by the decreased deposition of collagen in the IUA+DT group, as tested by immunostaining and Masson trichrome staining (Fig 6E). The pregnancy experiment showed that the number of live pups increased along with the decreased number of absorbed embryos (Fig 6F). Collectively, these observations supported that depletion of CD301b+ macrophages after establishing the IUA mouse model could effectively hinder the endometrial fibrosis and improve fertility.

**Figure 6.**
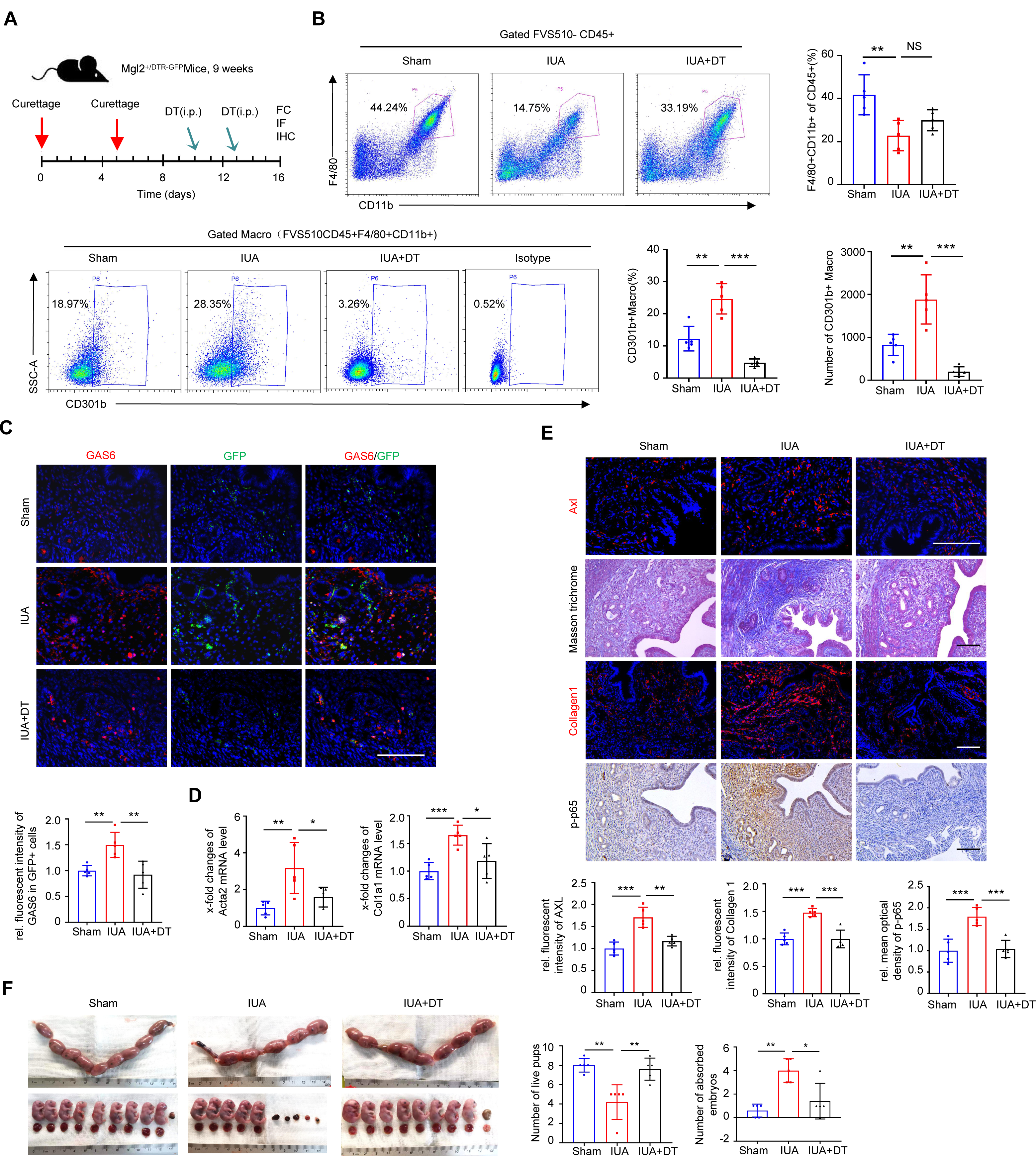
Depletion of CD301b+ macrophages in IUA mice reduces fibrosis burden and improves pregnancy outcome. A. Schema describing the approach to build the murine IUA model and delete CD301b+ macrophage by injecting diphtheria toxin (DT). B. Flow cytometric analysis of the proportions of CD11b+F4/80+ cells and CD301b+ macrophages in the uterus of Sham, IUA and IUA+DT mice (n=5 mice for each group). Macro, macrophage. C. Representative images of immunostaining for GAS6 (red) and GFP in the endometrium of Sham, IUA and IUA+DT mice (n=5 mice for each group). D. qRT-PCR analysis of Acta2 and Col1a1 expression in the uterus of Sham, IUA and IUA+DT mice (n=5 mice for each group). E. Representative images of immunostaining for AXL, Collagen1 and p-p65 and Masson trichrome staining in the endometria of Sham, IUA and IUA+DT mice (n=5 mice for each group). F. The number of live pups and absorbed embryos at 18.5 dpc among Sham, IUA and IUA+DT (n=5 mice for each group). Scale bar: 100μm. Data are presented as mean ± SEM. (B, C, D, E and F) One-way ANOVA with Tukey’s post hoc analysis. *, P<0.05; **, P<0.01; ***, P<0.001.

### Inhibition of AXL ameliorates endometrial fibrosis and restores endometrial function

To assess the efficacy of pharmacologic inhibition of the AXL pathway on endometrial fibrosis, a selective small molecule inhibitor against AXL, Bemcentinib, which has been used to perform clinical trials for treating cancers (Yule, et al.,2018; Bumm, et al.,2020), was given by oral administration (80 mg/kg, p.o., MedChemExpress) (Holland, et al.,2010; Lijnen, et al.,2011) in an IUA mouse model. We performed two kinds of time points for the treatment, one begun seven days later after the first curettage and one begun at the same day of the first curettage (Fig 7A, Appendix Fig S8A). The result showed that Bemcentinib inhibited the expression of Acta2 and Col1a1 expression at the mRNA level in the murine uterus by these two kinds of time points (Fig 7B, Appendix Fig S8B). Immunostaining showed that the expression levels of AXL, Collagen 1 and phospho-p65 were reduced and collagen deposition was diminished in the IUA mouse model by both kinds of treatment (Fig 7C, Appendix Fig S8C). The pregnancy experiment showed that the number of live pups was significantly increased and that of absorbed embryos was decreased in the IUA+BEM group by both kinds of treatment (Fig 7D, Appendix Fig S8D). These results indicated that administrating Bemcentinib after two curettages or prophylactically administrating Bemcentinib at the beginning of curettage were both effective and meliorated pregnancy outcome.

**Figure 7.**
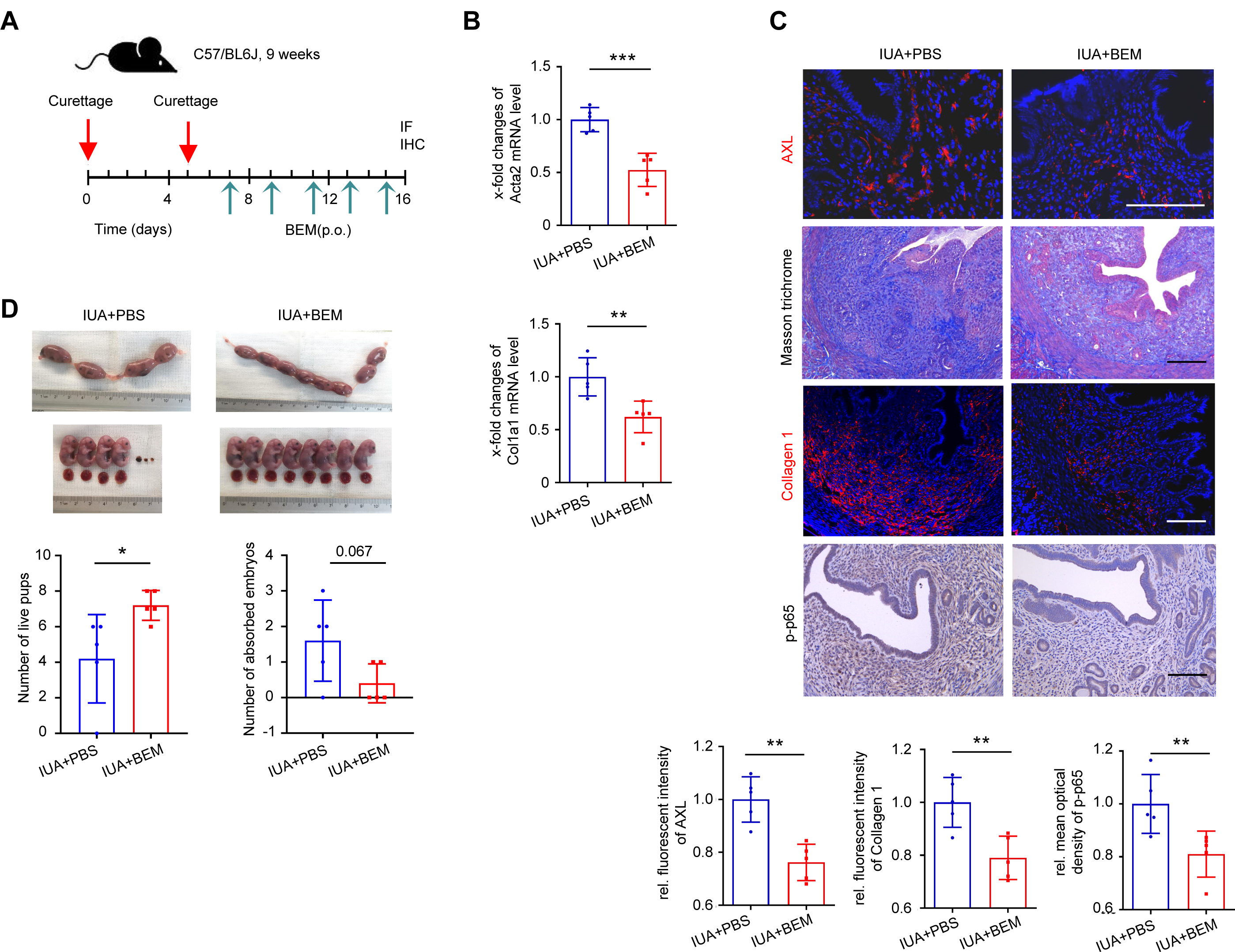
Bemcentinib mitigates established endometrial fibrosis and improves pregnancy outcome. A. Outline of the design of therapeutic dosing of Bemcentinib (BEM) in the established endometrial fibrosis. B. qRT-PCR analysis of Acta2 and Col1a1 expression in the uterus of IUA+PBS and IUA+BEM mice (n=5 mice for each group). C. Representative images of immunostaining for AXL, Collagen1, p-p65 and Masson trichrome staining in the endometrium of IUA+PBS group and IUA+BEM group (n=5 mice for each group). D. The number of live pups and absorbed embryos at 18.5 dpc between IUA+PBS group and IUA+BEM group (n=5 mice for each group). Scale bar, 100μm. Data are presented as mean ± SEM. (B, C for AXL and Collagen 1 and D) Two-tailed Student’s t-test. (C for p-p65) Mann-Whitney U test. *, P<0.05; **, P<0.01; ***, P<0.001.

## Discussion

In the present study, we focused on the relationship between the changes of macrophages and the formation of endometrial fibrosis in IUA. By analyzing the scRNA-seq datasets of the human IUA endometrium and experimental verification, we defined a critical profibrotic population, CD301+ macrophages that was required for the pathology of IUA and found its mechanism to function via the GAS6/AXL/NF-κB pathway. Furthermore, we demonstrated the effective targets of deletion of CD301+ macrophages or inhibition of AXL signaling activation by Bemcentinib to treat the endometrial fibrosis.

Endometrium is a dynamic changing tissue with the menstrual cycle and pregnancy and the number and subtypes of macrophages may also vary a lot at different phases. To avoid the physiological interference of menstrual cycle and pregnancy, we chose the endometrial biopsies at late proliferative phase which has a relatively stable state for the study to focus on the macrophages and pro-fibrotic cells in the endometrium. Few studies on the subtypes of endometrial macrophages at this phase were reported and the mechanisms of macrophages for fibrotic formation in IUA endometrium remain elusive. There were some subtypes of macrophages in other organs that have been recently reported to be related to fibrosis. For example, Lyve1^hi^MHCII^low^ macrophages, which were often closely associated with blood vessels across tissues, restrained the excessive leakage of inflammatory cells and overdeposition of collagen by regulating the vascular permeability (Chakarov, et al.,2019). GATA6+ macrophages located in the peritoneal cavity can rapidly aggregate to seal injuries and promote the repair of focal lesions, but they also formed extensive aggregates to promote the growth of intra-abdominal tissue known as peritoneal adhesions in iatrogenic surgical situations(Deniset, et al.,2019; Zindel, et al.,2021). CD301 was first studied in dendritic cells whose subset was required for the development of Th2 cells in dermis and submucosa(Kumamoto, et al.,2013), and then CD301b+ mononuclear phagocytes were found to maintain glucose metabolism and net energy balance through the secretion of resistin-like molecule alpha(Kumamoto, et al.,2016). Recently, it was reported that CD301+ macrophages have an important but paradoxical role in repair and fibrosis because in skin, it is critical to activate the proliferation of adipocyte precursors which promotes tissue repair during wound healing(Shook, et al.,2018); however, in heart tissues, CD301+ macrophages aggravate autoimmune cardiac valve inflammation and fibrosis(Meier, et al.,2018). This indicated that the role of CD301+ macrophages modified their role depending on their local microenvironment and accordingly exhibited high degrees of transcriptional and phenotypic heterogeneity and tissue-specific function. In this study, the single-cell RNA-seq analysis showed that CD301+ macrophages sent more signaling to myofibroblast-like cell and pseudo-time analysis predicted that this group of myofibroblast-like cells was differentiated from stromal cells. Our experiment results confirmed that this type of macrophage promoted stomal cells differentiation to myofibroblasts. Therefore, we think the reason why CD301+ macrophages were more closely related to myofibroblast-like cells might be that CD301+ macrophages promoted stromal cells differentiation and also played an important role in maintaining the survival and proliferation of myofibroblast-like cells.

In exploring the mechanism of pathogenesis in IUA, we found that the differentiation of both stromal cells and epithelial cells represented the key parenchymal cells involved in IUA pathogenesis. The decreased expression of VEGF165 in the IUA endometrium would target stromal cells to differentiate into myofibroblasts by dampening the DLL4/Notch4/Smad7 axis(Lv, et al.,2019), and ΔNp63α, which is ectopically expressed in endometrial epithelial cells (EECs) of IUA patients and regulated by circPTPN12/miR-21-5p(Zhao, et al.,2017; Song, et al.,2021), could promote the expression of SNAI1 by the DUSP4/GSK3B pathway to induce EEC-EMT and endometrial fibrosis(Zhao, et al.,2020). In this study, we screened out GAS6, which was highly expressed in CD301+ macrophages, was a crucial secretory molecule that induces profound dysfunction of stromal cells via previously unappreciated AXL/NF-κB signaling in the endometrium. AXL has recently been reported to drive profibrotic phenotypes in fibroblasts or endothelial cells of renal and pulmonary fibrosis(Möller-Hackbarth, et al.,2021; Yang, et al.,2021) and we found that its inhibitor Bemcentinib could obviously restrain the progress of fibrosis and improve functional recovery in the endometrium of IUA.

Bemcentinib has now been used for clinical trials to treat acute myeloid leukemia (Sonja, et al.,2019), non-small alveolar lung cancer (Lauren, et al., 2021) and the common side effects are ECG QT prolongation, diarrhea and nausea which are manageable and/or reversible. The off-target effect of Bemcentinib *in vivo* has not been documented. In our study, we found that Bemcentinib administration can increase live pups and decrease absorbed embryos, at least in part, demonstrating that it may have minimal toxicity for reproduction at this dose. Further studies are needed to investigate the off-target effects of Bemcentinib *in vivo*. Some other studies have tried to use stem cells, M-CSF, bFGF, auto-crosslinked hyaluronic acid, etc., for treating endometrial fibrosis of IUA and its recurrence (Cao, et al.,2018; Jiang, et al.,2019; Liu, et al.,2020; Santamaria,2018), but our research is the first to demonstrate that the resolution and prevention of endometrial fibrosis can be accelerated by targeting a receptor tyrosine kinase in the endometrium, which is a pharmacological intervention.

In summary, our study unravels that CD301+ macrophages are increased in the endometria of IUA and they aggravate stromal cell differentiation into myofibroblasts via the GAS6/AXL/NF-κB axis to accelerate profibrotic responses during the progression of endometrial fibrosis in IUAs. Treatments aimed at fine-tuning the number of CD301+ macrophages or restraining its downstream molecules could be of good clinical value.

## Materials and Methods

### Human samples

The human endometrial samples and procedures involved in this study were approved by the Ethics Committee of Nanjing Drum Tower Hospital, The Affiliated Hospital of Nanjing University Medical School. Human endometrial samples were collected at the late proliferative phase of the menstrual cycle from women aged 22-40 years during hysteroscopic examination at the Infertility Consulting Clinic at Nanjing Drum Tower Hospital. The late proliferative phase was defined based on follicle size between 15 and 18 mm by ultrasonography and a low level of serum progesterone. Samples were taken from two sites in the uterine body or fundus of IUA patients whose endometria were diagnosed with scores > 8 based on criteria recommended by the American Fertility Society (American Fertility Society,1988), and non-IUA patients who had tubal infertility with normal ovary function, normal menstrual blood volume, and endometrium thickness ≥ 7 mm immediately before ovulation. Women were excluded from the tissue collection if they were under any of the following condition: positive serological tests for human immunodeficiency virus, hepatitis B/C virus, syphilis; history of tuberculosis; chronic endometritis; vaginal bacterial or fungal. Patients with uterine malformations determined by ultrasonography have been ruled out before sample collection.

To explore the changes of cell population ratios, differential gene expression profiles and the cell-cell communications in the endometria between IUA patients and normal controls, we randomly selected three endometrial samples from the initial ten IUA patients and three endometrium samples from the initial ten controls to perform single-cell RNA sequencing between September 2019 and December 2019. After obtaining the results of single-cell RNA sequencing, we further enrolled additional 18 IUA patients and 26 controls until December 2021. Thus, a total of 28 IUA patients and 36 controls were included in the present study. To validate the results of single-cell RNA sequencing, we randomly selected 20 endometrium samples from each group to conduct flow cytometry; five samples from each group were used for immunostaining; three and five samples from the remaining samples of the control group, were used for CD301-/+macrophage sorting and isolation of the primary stromal cells for in vitro study, respectively. The clinical information of all patients and controls for single-cell RNA sequencing was listed in Appendix Table S1, and for other experiments, it was shown in Appendix Table S4.

### Animals

All animal experiments were carried out in accordance with the guidelines of the Experimental Animals Management Committee (Jiangsu Province, China) and were approved by the Ethics Review Board for Animal Studies of Nanjing Drum Tower Hospital. Nine to ten-week-old C57BL/6 female virgin mice, weighing 18-20 g, were purchased from the Experimental Animal Center of Nanjing Medical University (Nanjing, China). To generate Mgl2+/DTR-GFP (Mgl2-DTR) mice, we obtained help from GemPharmatech Co., Ltd. to construct the CRISPR/Cas9 system targeting the mouse Mgl2 gene and insert the donor vector, DTR-GFP-PolyA into the translation start codon site of the mouse Mgl2 gene to conditionally abolish the mouse endogenous Mgl2 gene. The CRISPR/Cas9 system was constructed in vitro and then microinjected into the fertilized eggs of C57BL/6JGpt mice to obtain positive F0 generation mice. The F0-positive mice were mated with C57BL/6JGpt mice, and their pups were genotyped by PCR, followed by sequence analysis. GFP fluorescence in mice confirmed the selective expression of the DTR-GFP fusion protein in CD301b+ cells. To deplete CD301b-expressing cells, 100 μL of 10 μg/mL diphtheria toxin (DT) was administered intraperitoneally to Mgl2-DTR mice, and the depletion lasted for at least 6 days after a single DT injection (Appendix Fig S6).

### Single cell RNA-seq data processing

The endometrium samples were rinsed several times with PBS until there were no obvious blood clots, cut into small pieces, digested with 0.1% trypsin (Wisent Inc., Canada) for 8 min and then transferred into 0.8 mg/mL Collagenase Type I (Sigma-Aldrich, USA) for 60 min at 37 °C and 5% CO2 with regular vigorous shaking in a humidified incubator. The released endometrial cells were filtered through a 70μm cell strainer (BD Biosciences, USA), centrifuged, and resuspended in 5mL of red blood cell lysis buffer (eBiosciences, USA) for 8 min to exclude any remaining red blood cells. Then they were resuspended in PBS and used for single-cell 3’-cDNA library preparation followed by the 10X Genomics Chromium Single Cell 3’ Reagent Kit protocol. Single-cell encapsulation, complementary DNA (cDNA) library synthesis and RNA-sequencing were completed by Gene Denovo (Guangzhou, China).

Single cell libraries were sequenced on Illumina NovaSeq instruments using 150 nt paired-end sequencing. Reads were processed using the Cell Ranger 4.0.0 pipeline with the default and recommended parameters. FASTQ files generated from Illumina sequencing output were aligned to the human reference genome (GRCh38) using the STAR algorithm. The FASTQ data of normal controls were also used in our published study (Lv, et al.,2022). The unique molecular identifier (UMI) count matrix was then transformed into the Seurat objects using the R package Seurat v2(Butler, et al.,2018). The raw gene expression measurements for each cell were normalized by dividing them by the total expression followed by scale factor-multiplying (x 10,000) and log-transformation ln (UMI-per-10,000+1) in the Seurat toolkit. Cells that expressed fewer than 200 genes or mitochondrial gene content > 15% of the total UMI count were excluded.

### Dimension reduction and identification of major cell clusters

To integrate and correct the data from different batches, canonical correspondence analysis (CCA)(Butler, et al.,2018) was performed which removed technical variation and identified common cell types and markers. The normalized gene expression matrix generated after data preprocessing was used to identify the major cell clusters by applying dimension reduction and clustering. Uniform Manifold Approximation and Projection (UMAP) method have been used for visualization of unsupervised clustering at the resolution of 0.5. Detailed descriptions of the cellular subsets and their marker genes were included in the figures and main text of the relevant sections. The cell type of each clustered was determined by known markers of individual cell types combined with differentially expressed genes in the cluster compared to all other cells.

### CellPhoneDB

To investigate potential interactions across cellular subsets, cell-cell communication analysis was performed using CellPhoneDB, which was a publicly available repository of curated receptors and ligands and their interactions that enabled a comprehensive, systematic analysis of cell-cell communication molecules(Efremova, et al.,2020). Enriched receptor-ligand interactions between two cell types were derived on the basis of expression of a receptor by one cell type and a ligand by another cell type. The member of the complex with the minimum average expression was considered for the subsequent statistical analysis. A null distribution of the mean of the average ligand and receptor expression in the interacting clusters was generated by randomly permuting the cluster labels of all cells. The P value for the likelihood of cell-type specificity of a given receptor-ligand complex was calculated on the basis of the proportion of the means that were as high as or higher than the actual mean. A P value < 0.05 was considered statistically significant. Heatmaps were generated based on the number of ligand-receptor pairs that were statistically significant in cell-to-cell interactions. The cells on the Y-axis were the ones expressing legends, and the cells on the X-axis were the ones expressing receptors. The color of the heatmaps was based on the number of ligand-receptor pairs that were statistically significant in cell-to-cell interactions.

### Pseuduotime trajectory analysis

The single cell trajectory was analyzed using a matrix of cells and gene expression by Monocle (Version 2.6.4). Monocle reduced the space down to one with two dimensions and ordered the cells (sigma = 0.001, lambda = NULL, param.gamma = 10, tol = 0.001). Once the cells were ordered, we could visualize the trajectory in the reduced dimensional space. The trajectory had a tree-like structure, including tips and branches. Often, single-cell trajectories included branches. The branches occurred because cells executed alternative gene expression programs. Branches appeared in trajectories during development, when cells made fate choices: one developmental lineage proceeded down one path, while the other lineage produced a second path. Monocle developed BEAM to test for branch-dependent gene expression by formulating the problem as a contrast between two negative binomial GLMs.

### Flow cytometry

The method of human endometrium or mouse uterine being dissected and digested into single cell was the same as that for single-cell RNA sequencing above. Then the released cells were resuspended in FACS staining buffer (0.1% BSA in PBS). To examine immune cell subsets, cells were stained for viability (FVS510) and the indicated antibodies for 15 min. Data were acquired using CytoFLEX (Beckman Coulter, USA) and analysis was performed using CytExpert v2.3 (Beckman Coulter, USA). The antibodies used were shown in Appendix Table S6.

### Magnetic separation and fluorescence-activated cell sorting

The method for human endometrium to be dissected and digested into single cells was the same as above. Cells were magnetically labeled and separated according to the instructions of CD45 MicroBeads Human (order no. 130-045-801, Miltenyi Biotec). Then, CD45+ endometrial cells were collected and stained with the following antibodies for 15 minutes: CD14-FITC, CD301-APC and FVS510. CD301+ macrophages and CD301-macrophages were sorted into RPMI 1640 (Sigma) with 10% fetal bovine serum (FBS; Gibco) for culture.

### Cell culture

Human endometrial stromal cells (hESCs) from normal endometrial biopsies were dissected and digested according to the method described above and then filtered through a 40 μm sieve gauze to separate the stromal cells from the glands. Cells were inoculated in DMEM/F12 (Wisent Inc) containing 10% FBS (Gibco) and cultured at 37 ℃ with 5% CO_2_ and saturated humidity. The hESCs from passages 2-4 were used for all experiments. Chemicals and recombinant proteins were used in the article are recombinant human Gas6 (order no. 885-GSB), Bemcentinib (order no. HY-15150), JSH23 (order no. HY-13982).

### Quantitative real-time PCR

Total RNA from the tissues or cultured cells was extracted from TRIzol Universal Reagent (TIANGEN). One microgram of RNA was reverse transcribed into cDNA using HiScript^®^ III RT Super Mix for qPCR (+gDNA wiper) (Vazyme). Quantitative real-time PCR (qRT-PCR) was performed as previously described(Lv, et al.,2019). Differences among the target gene expression levels were estimated by the ΔΔ*C*_t_ method and normalized to the level of 18S. The values were the mean ± SEM. The primers used in this study were listed in Appendix Table S5.

### Immunohistochemistry

Uterine tissues were fixed with 4% phosphate-buffered paraformaldehyde overnight and then dehydrated and embedded in paraffin. Paraffin sections (2 μm) were prepared, dewaxed, hydrated, and the endogenous peroxides were quenched with 3% H_2_O_2_. After heat-mediated antigen retrieval, the slides were incubated with primary antibodies for 2 h at 37 ℃. After incubation with HRP-conjugated secondary antibodies, the sections were exposed to DAB and counterstained with hematoxylin. A Leica DM6B microscope was used for visualization. Slides were observed by two pathologists independently in a double-blind fashion. Immunohistochemical staining was quantified by mean optical density using Image-Pro Plus v6. The antibodies used were shown in Appendix Table S6.

### Immunofluorescence

Tissues were fixed with 4% phosphate-buffered paraformaldehyde overnight, protected in graded sucrose, embedded in OCT and finally frozen. The cryostat sections were incubated in blocking buffer (1% bovine serum albumin, 0.1% Triton-X in PBS) at room temperature for 30 minutes. Primary and secondary antibodies were added, followed by staining with DAPI. The staining was viewed using a Leica DM6B microscope by two pathologists independently in a double-blind fashion. The results were quantified by the mean fluorescence intensity or the number of positive cells using Image-Pro Plus v6. The antibodies used were shown in Appendix Table S6.

### Western blot

After examining the protein concentration of the cell lysates, each sample with equal amounts of protein was subjected to SDS–PAGE and transferred to a PVDF membrane (Millipore, Burlington, MA). The membranes were incubated with the primary antibody at 4 ℃ overnight, followed by HRP-conjugated secondary antibodies at room temperature for 1 hour. The signals were visualized with ECL solution (Fdbioscience), and the protein expression levels were quantified by analysis of the integrated density normalized to the level of β-actin using ImageJ. The antibodies used were shown in Appendix Table S6.

### Bulk RNA-sequencing and analysis

hESCs were isolated from normal human normal endometrium and treated with 50 ng/mL GAS6 for 72 h. Control hESCs were treated with the same volume of PBS. Total RNA was extracted using TRIzol reagent. Transcriptome sequencing analysis was performed on an Illumina NovaSeq6000 by Gene Denovo Biotechnology Co. (Guangzhou, China). Differential expression analysis between two different groups was performed with DESeq2 software. Genes with a false discovery rate (FDR) below 0.05 and absolute fold change≥1.5 were considered differentially expressed genes (DEGs). KEGG which is the major public pathway-related database for pathway enrichment analysis was performed for all DEGs. It identifies significantly enriched metabolic pathways or signal transduction pathways in DEGs comparing with the whole genome background. Scatterplot figure was made for the top 10 enriched pathways.

The published dataset GSE105789 for gene expression between CD301b+ macrophage and F4/80-immune cells was extracted for differential expression analysis by DESeq2 software. Genes with a false discovery rate (FDR) below 0.05 and absolute fold change≥1.5 were considered differentially expressed genes (DEGs). A volcano plots was made to show statistical significance (P value) versus magnitude of change (fold change).

### Animal model

The IUA model was built using mice that were 9-10 weeks of age, 18-20 g and at estrum, which corresponds to the late-proliferating phase in humans, defined by vaginal smears. All the mice were anesthetized with isoflurane. To generate the IUA model, laparotomy was performed to expose the uterus, and the cervix was poked to allow a rough-surfaced needle to be inserted all the way through the lumen, which was then scratched up and down 100 times until the walls of the uterine cavity became rough. The lumen was scratched again five days later. Control sham mice underwent the same procedure without scratching. Uterine tissues were collected at estrum, around 16 days after the first surgery. The other mice used to assess pregnancy outcomes began to be mated with wild-type male mice at the 13th day after first surgery and Day 0.5 of pregnancy (0.5dpc) was defined as the day when vaginal plugs were recognized.

### Statistical analysis

Data analysis was performed using GraphPad Prism 9. Normal distribution data were analyzed by unpaired t test for comparisons between two groups or by one-way ANOVA with Tukey’s post-hoc test for multiple comparisons. A nonparametric test was used when data were not normally distributed. Data were presented as mean ± SEM. A P value < 0.05 was considered statistically significant. In all figures, one, two and three asterisks indicated *P < 0.05, **P < 0.01 and ***P < 0.001, respectively.

## Data availability

The sequencing datasets are available in the NCBI Gene Expression Omnibus and Sequence Read Archive: ID: PRJNA730360 for scRNA-seq of normal endometria samples, PRJNA788201 for scRNA-seq of IUA endometria samples, PRJNA788544 for bulk RNA-seq of hESCs treated by GAS6. All other data generated or analyzed during this study are included within the paper. All reagents used in this study are commercially available.

## Acknowledgements

This work was supported by National Key Natural Science Foundation of China (82030040), The Strategic Priority Research Program of the Chinese Academy of Sciences (XDA16040300), National Natural Science Foundation of China (82071600, 82171618, 81971336), National Key R&D Program of China (2021YFC2701603) and Jiangsu Biobank of Clinical Resources (BM2015004).

## Author contributions

**Haining Lv:** Conceptualization; formal analysis; investigation; methodology; writing-original draft; writing-review and editing; **Haixiang Sun**: Conceptualization; supervision; funding acquisition; writing-review and editing. **Limin Wang:** Formal analysis, investigation; methodology; writing-review and editing. **Simin Yao:** Investigation; methodology; writing-review and editing. **Dan Liu:** Investigation; methodology; writing-review and editing. **Xiwen Zhang:** Investigation; methodology; writing-review and editing. **Zhongrui Pei:** Investigation; methodology; writing-review and editing. **Jianjun Zhou:** Investigation; methodology; writing-review and editing. **Huiyan Wang:** Investigation; methodology; writing-review and editing. **Jianwu Dai:** Investigation; methodology; writing-review and editing. **Guijun Yan:** Investigation; methodology; writing-review and editing. **Lijun Ding:** Investigation; methodology; writing-review and editing. **Zhiyin Wang:** Investigation; methodology; writing-review and editing. **Chenrui Cao:** Investigation; methodology; writing-review and editing. **Guangfeng Zhao:** Conceptualization; supervision; funding acquisition; writing-original draft; writing-review and editing. **Yali Hu:** Conceptualization; supervision; funding acquisition; writing-original draft; writing-review and editing.

## The Paper Explained Problem

Intrauterine adhesion (IUA) characterized by endometrial fibrosis, is a main disease of uterine infertility, however access to effective treatment is a big challenge due to its high recurrence rate after treatment and the rudimentary understanding of the pathogenesis.

## Results

We captured CD301+ macrophage that has an important role in the progression of IUA by single-cell RNA sequencing and validated that CD301+ macrophage is increased in endometrium of IUA which promotes endometrial stromal cells to differentiate into myofibroblasts and aggravates endometrial fibrosis. Mechanistically, CD301+ macrophage secretes GAS6 to bind its receptor AXL and then activates NF--κB pathway to up-regulate the profibrotic protein synthesis. Targeted deletion of CD301+ macrophage or pharmacological inhibitor of AXL prevents endometrial fibrosis and improves the pregnancy outcome in mice.

## Impact

Our findings demonstrate the therapeutic potential of macrophage-modulation to suppress endometrial fibrosis and increase the success in pregnancy.

