## Supplementary Figures and for "Targeting CD301+ macrophage inhibits endometrial fibrosis and improves pregnancy outcome"

**Supplementary figures: 8**

**Supplementary tables: 6**

**Appendix Figure S1**

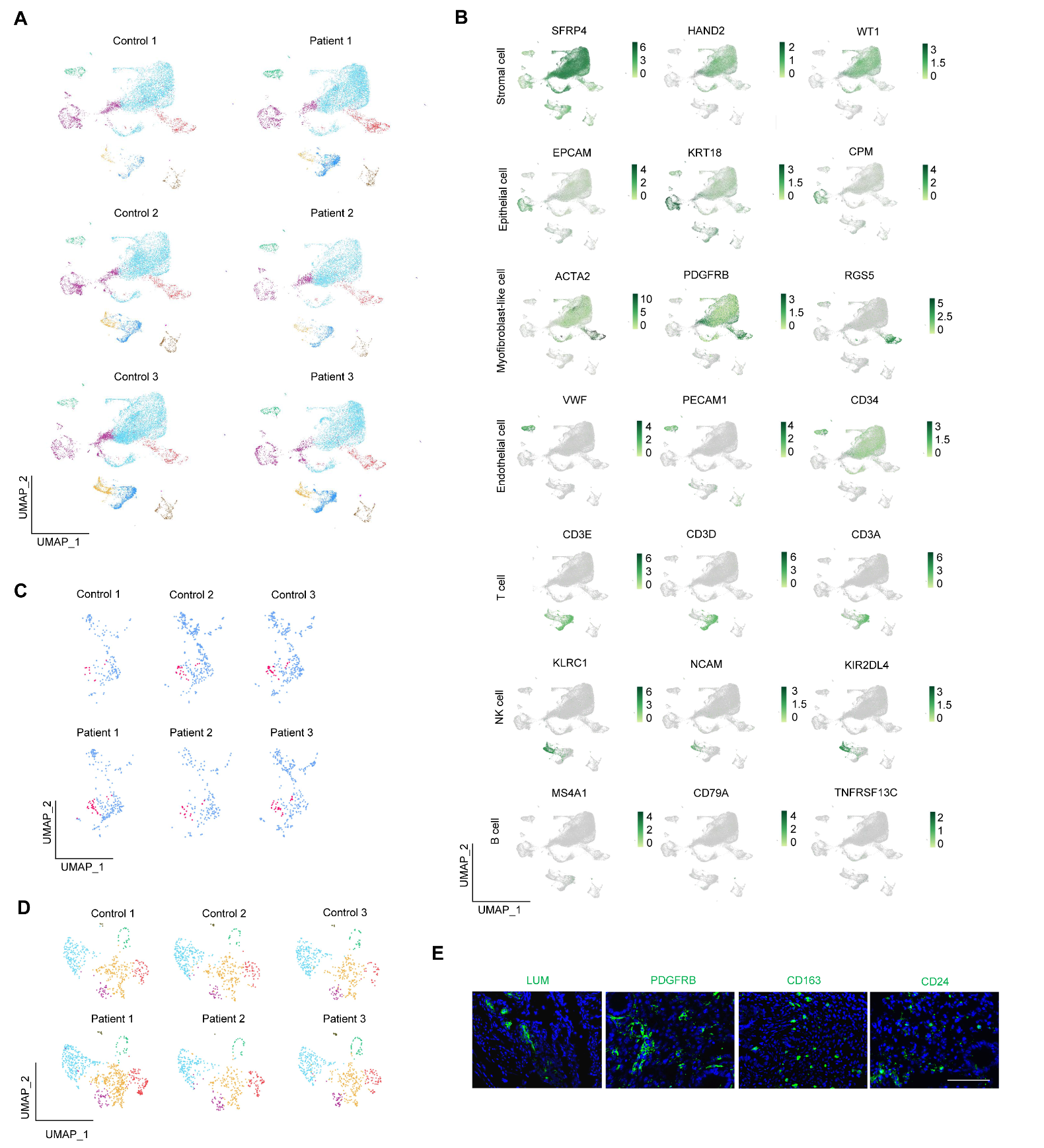

**Appendix Figure S1 Endometrial cells atlas**.

A. The distribution of each type of cells by UMAP in the controls (n=3 samples) and IUA patients (n=3 samples).
B. The distribution of marker genes for each type of cells (n=6 samples).
C. The distribution of dendritic cells and macrophage by UMAP from controls (n=3 samples) and IUA patients (n=3 samples).
D. The distribution of subtypes of myofibroblast-like cells by UMAP in controls (n=3 samples) and IUA patients (n=3 samples).
E. Localization of LUM, PDGFRB, CD163, and CD24 in endometrium of normal controls examined by immunofluorescence. Scale bar: 100μm.

**Appendix Figure S2**

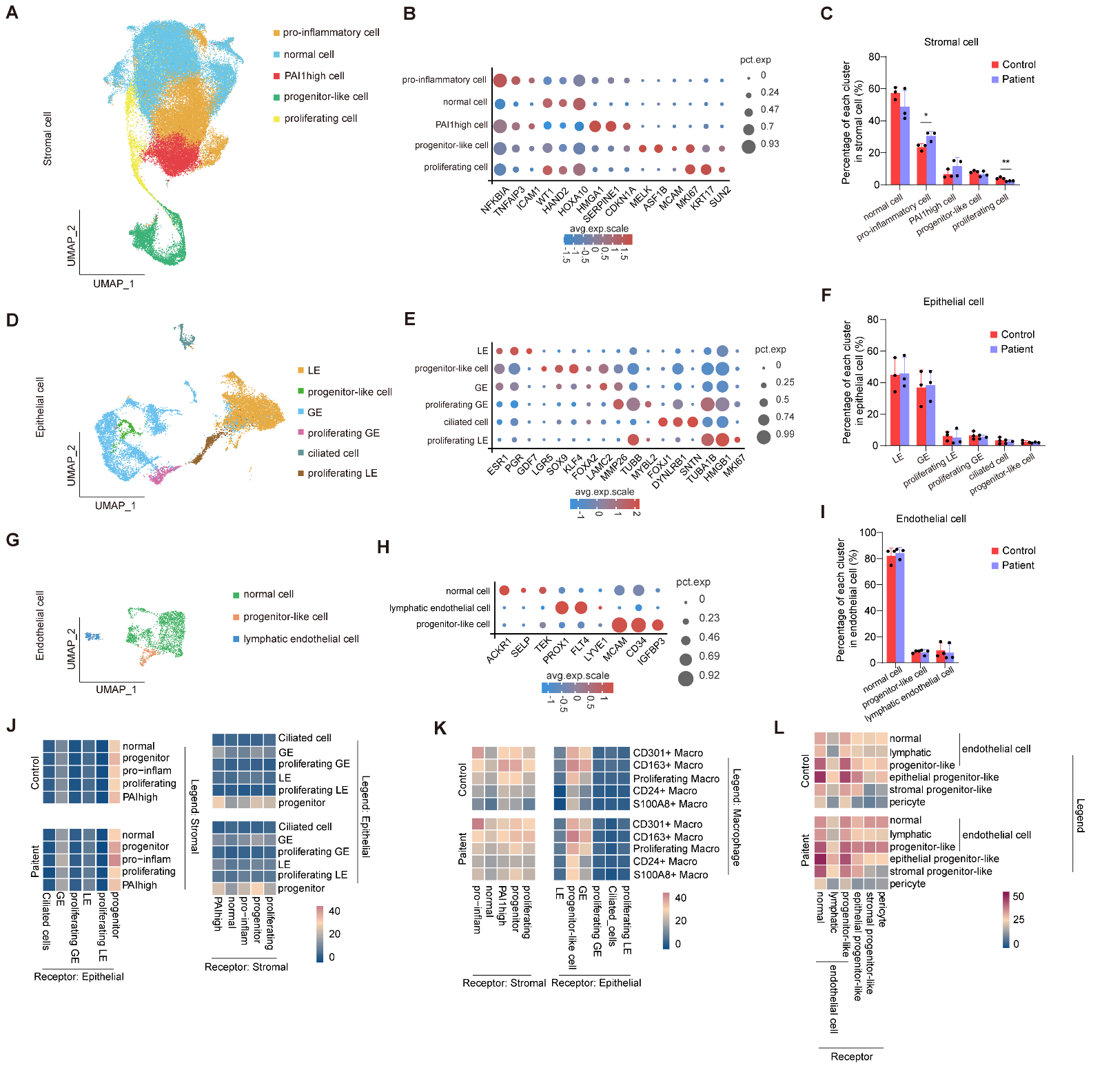

**Appendix Figure S2 Characteristics of subpopulations of stromal cells, epithelial cells and endothelial cells.**

A. The UMAP plot embedding subpopulations of stromal cells annotated by cell type/state from controls and patients with IUA (n=6 samples).
B. Bubble diagram showing the expression of marker genes for each cell subtype of stromal cells (n=6 samples).
C. The percentage of each cell subtype in total stromal cells between controls and patients with IUA (n=3 samples for each group).
D. The UMAP plot embedding subpopulations of epithelial cells annotated by cell type/state from controls and patients with IUA (n=6 samples).
E. Bubble diagram showing the expression of marker genes for each cell subtype of epithelial cells (n=6 samples).
F. The percentage of each cell subtype in total epithelial cells between controls and patients with IUA (n=3 samples for each group).
G. The UMAP plot embedding subpopulations of endothelial cells annotated by cell type/state from controls and patients with IUA (n=6 samples).
H. Bubble diagram showing the expression of marker genes for each cell subtype of endothelial cells (n=6 samples).
I. The percentage of each cell subtype in total endothelial cells between controls and patients with IUA (n=3 samples for each group).
J. The abundance of connection between subpopulation of stromal cells and epithelial cells utilizing CellPhoneDB in controls and patients with IUA (n=3 samples for each group).
K. The abundance of connection between subpopulation of macrophages (as legends’ cells) and subpopulation of stromal cells or epithelial cells (as receptors’ cells) utilizing CellPhoneDB in controls and patients with IUA (n=3 samples for each group).
L. The abundance of connection among subpopulation of endothelial cells, epithelial progenitor-like cells, stromal progenitor-like cells, and pericyte utilizing CellPhoneDB in controls and patients with IUA (n=3 samples for each group).

Data are presented as mean ±SEM. (C, F and I) Two-tailed Student’s t-test. *, P<0.05; **, P<0.01.

**Appendix Figure S3**

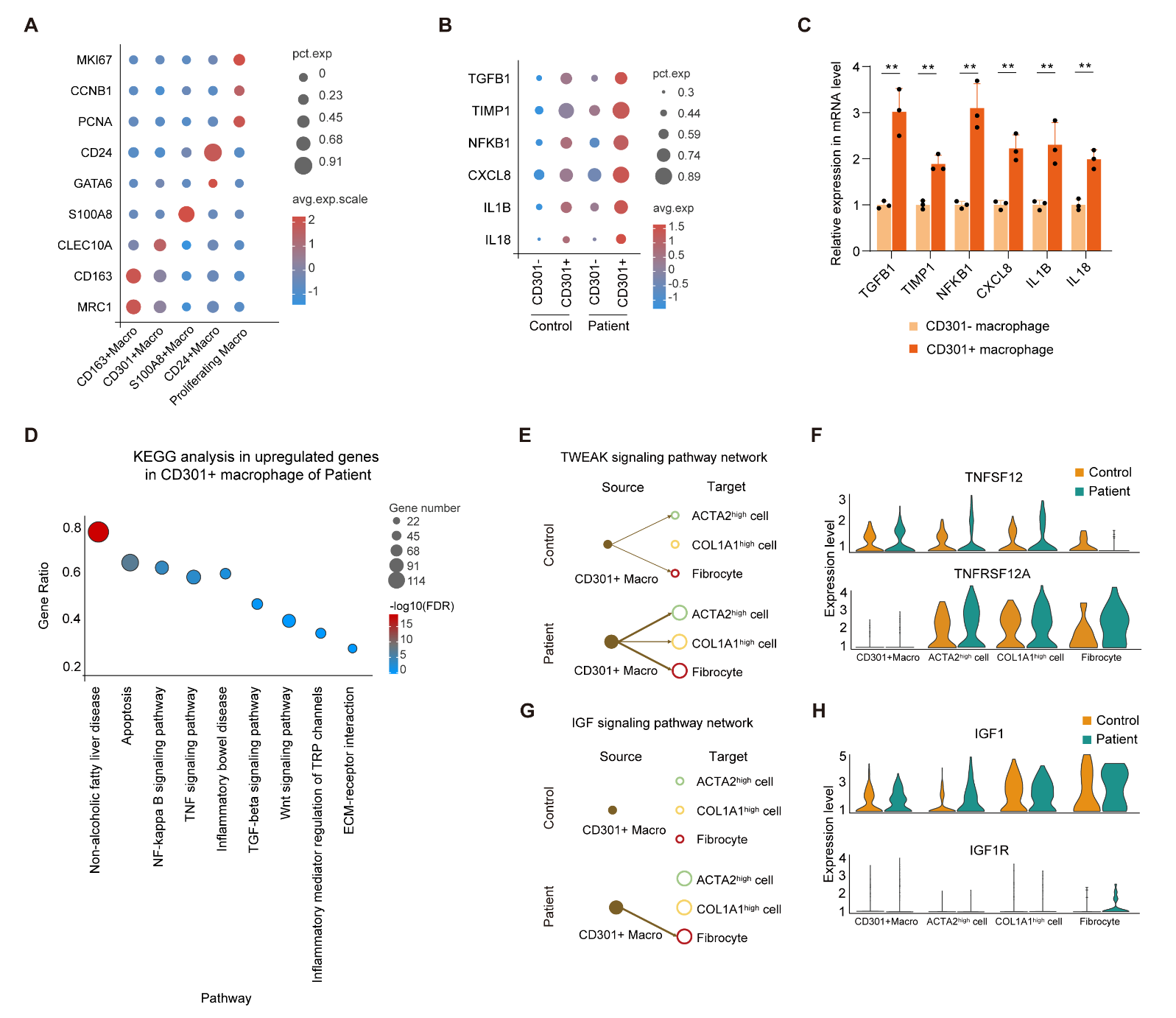

**Appendix Figure S3 Characteristics of macrophage and the communication between CD301+ macrophage and myofibroblast-like cells**

A. Bubble diagram showing the expression of marker genes for each subpopulation of macrophages (n=6 samples).
B. Bubble diagram showing the genes related to inflammation which are significantly high-expressed in CD301+ macrophage (n=3 samples for each group).
C. qRT-PCR analysis of indicated gene expression in sorted endometrial CD301- macrophage and CD301+ macrophage (n=3 biological replicates for each group).
D. KEGG analysis in significantly upregulated genes in CD301+ macrophage of patients compared to controls using scRNA-seq data (n=3 samples for each group).
E. Hierarchy diagram showing the interaction from CD301+ macrophage to pro-fibrotic subpopulation of myofibroblast-like cells in TWEAK signaling pathway network analyzed by CellChat (n=3 samples for each group). The size of circle diameter represents relative number of cells and the size of line represents relative strength of subpopulation interaction between controls and patients
F. Gene expression patterns of TWEAK signaling pathway in CD301+ macrophage, ACTA2^high^ cell, COL1A1^high^ cell and fibrocyte analyzed by CellChat (n=3 samples for each group).
G. Hierarchy diagram showing the interaction from CD301+ macrophage to pro-fibrotic subpopulation of myofibroblast-like cells in IGF signaling pathway network analyzed by CellChat (n=3 samples for each group). The size of circle diameter represents relative number of cells and the size of line represents relative strength of subpopulation interaction between controls and patients.
H. Gene expression patterns of IGF signaling pathway in CD301+ macrophage, ACTA2^high^ cell, COL1A1^high^ cell and fibrocyte analyzed by CellChat (n=3 samples for each group).

Data are presented as mean ±SEM. (C) Two-tailed Student’s t-test. **, P<0.01.

**Appendix Figure S4**

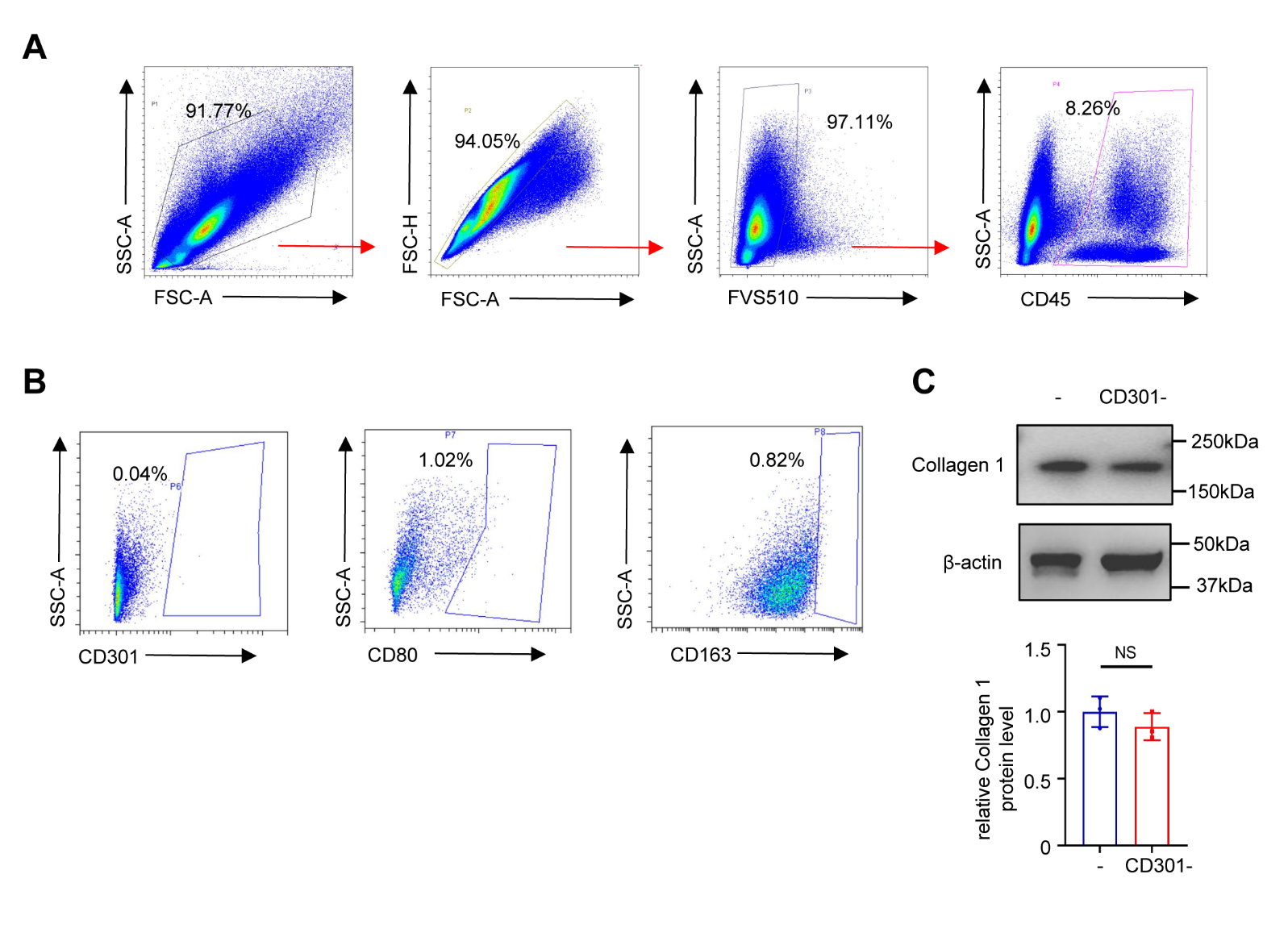

**Appendix Figure S4 Flow cytometric analysis of human endometrial tissue and the effect of CD301- macrophage on stromal cells.**

A. Flow cytometry gating strategy for FVS510-CD45+ cells.
B. Flow cytometric analysis of isotype of CD301, CD80 and CD163 expression in endometrium.
C. Western blot analysis of Collagen 1 expression in hESCs only and hESCs treated supernatants of CD301-macrophage for 48h (n=3 technical replicates for each group).

Data are presented as mean ± SEM. (C) Two-tailed Student’s t-test. NS, not significant.

**Appendix Figure S5**

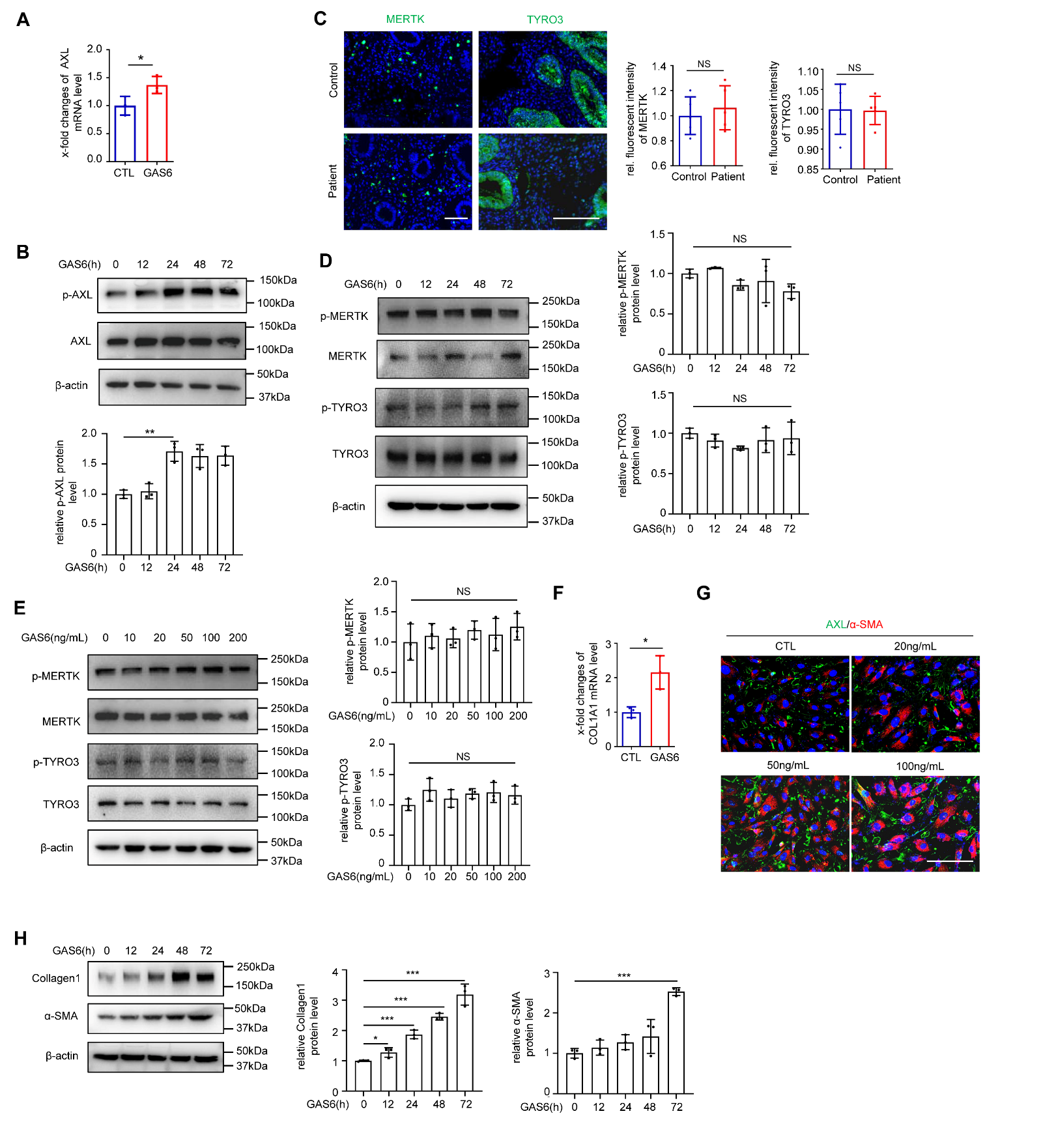

**Appendix Figure S5. Expression of MERTK and TYRO3 in human endometrial sections and cells.**

A. qRT-PCR analysis of AXL expression in hESCs treated with or without GAS6 (50ng/mL) for 72h (n=3 biological replicates for each group).
B. Western blot analysis of phospho-AXL (p-AXL) and AXL in hESCs treated with GAS6 (50ng/mL) at time of 12, 24, 48 and 72h (n=3 technical replicates for each group).
C. Localization of MERTK and TYRO3 in endometrium of normal controls and patients with IUA (n=5 samples for each group). Scale bar: 100μm.
D. Western blot analysis of phosphorylated MERTK, TYRO3 (pMERTK, pTYRO3) and MERTK, TYRO3 in hESCs treated with GAS6 (50ng/mL) at time of 12, 24, 48 and 72h (n=3 technical replicates for each group).
E. Western blot analysis of phosphorylated MERTK, TYRO3 (pMERTK, pTYRO3) and MERTK, TYRO3 in hESCs treated with GAS6 at dose of 10, 20, 50, 100 and 200ng/mL (n=3 technical replicates for each group).
F. The mRNA level of COL1A1 expression in hESCs after treatment with GAS6 (50ng/mL) for 72h analyzed by qRT-PCR (n=3 biological replicates for each group).
G. Localization of AXL and α-SMA in hESCs after treatment with GAS6 at dose of 20, 50 and 100ng/mL for 72h. Scale bar: 50μm.
H. Western blot analysis of α-SMA and Collagen1 in hESCs treated with GAS6 (50ng/mL) at time of 12, 24, 48 and 72h (n=3 technical replicates for each group).

Data are presented as mean ± SEM. (A, C and F) Two-tailed Student’s t-test. (B, D, E, H) One-way ANOVA with Tukey’s post hoc analysis. *, P<0.05; **, P<0.01; ***, P<0.001. NS, not significant.

**Appendix Figure S6**

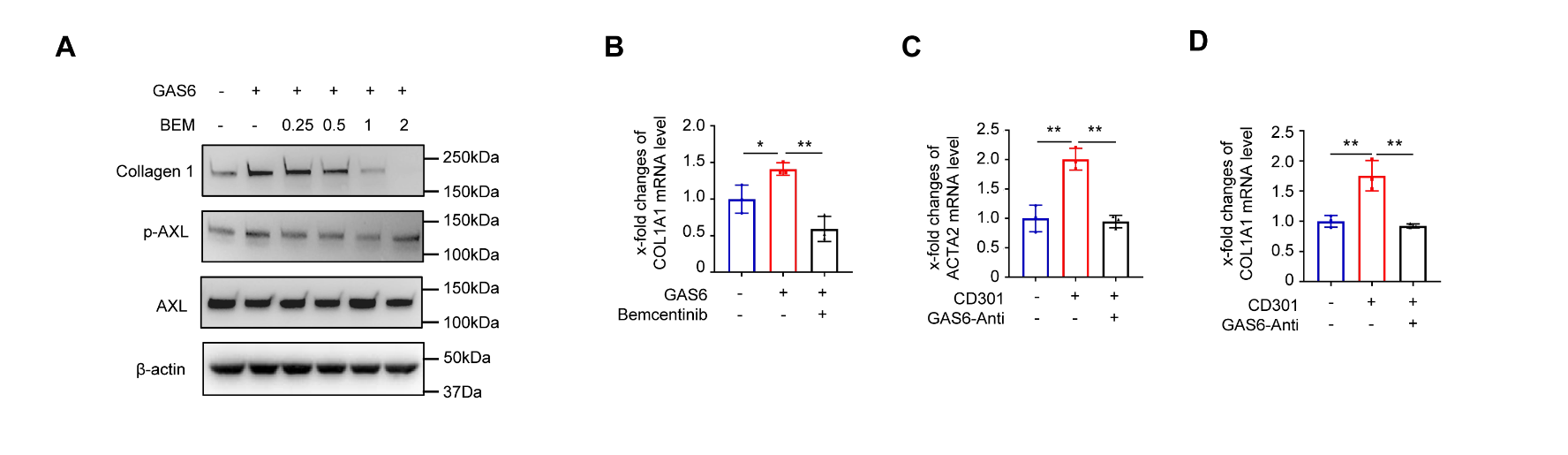

**Appendix Figure S6. Expression of downstream genes under the treatment of Bemcentinib or GAS6 antibody.**

A. Western blot analysis of Collagen 1, phosphorylated AXL and AXL in hESCs treated with Bemcentinib at the dose of 0, 0.25, 0.5, 1, 2μM for 1 hour, followed by GAS6 (50ng/mL) for 72h.
B. The mRNA level of COL1A1 expression in hESCs after treatment with Bemcentinib (1μM) for 1 hour, followed by GAS6 (50ng/mL) for 72h (n=3 biological replicates for each group).
C. qRT-PCR analysis of ACTA2 expression in hESCs treated with the supernatants of CD301+ and CD301- macrophage and GAS6 neutralizing antibody (2.5μg/mL) for 72h (n=3 biological replicates for each group).
D. qRT-PCR analysis of COL1A1 expression in hESCs treated with the supernatants of CD301+ and CD301- macrophage and GAS6 neutralizing antibody (2.5μg/mL) for 72h (n=3 biological replicates for each group).

Data are presented as mean ± SEM. (B-D) One-way ANOVA with Tukey’s post hoc analysis. *, P<0.05; **, P<0.01.

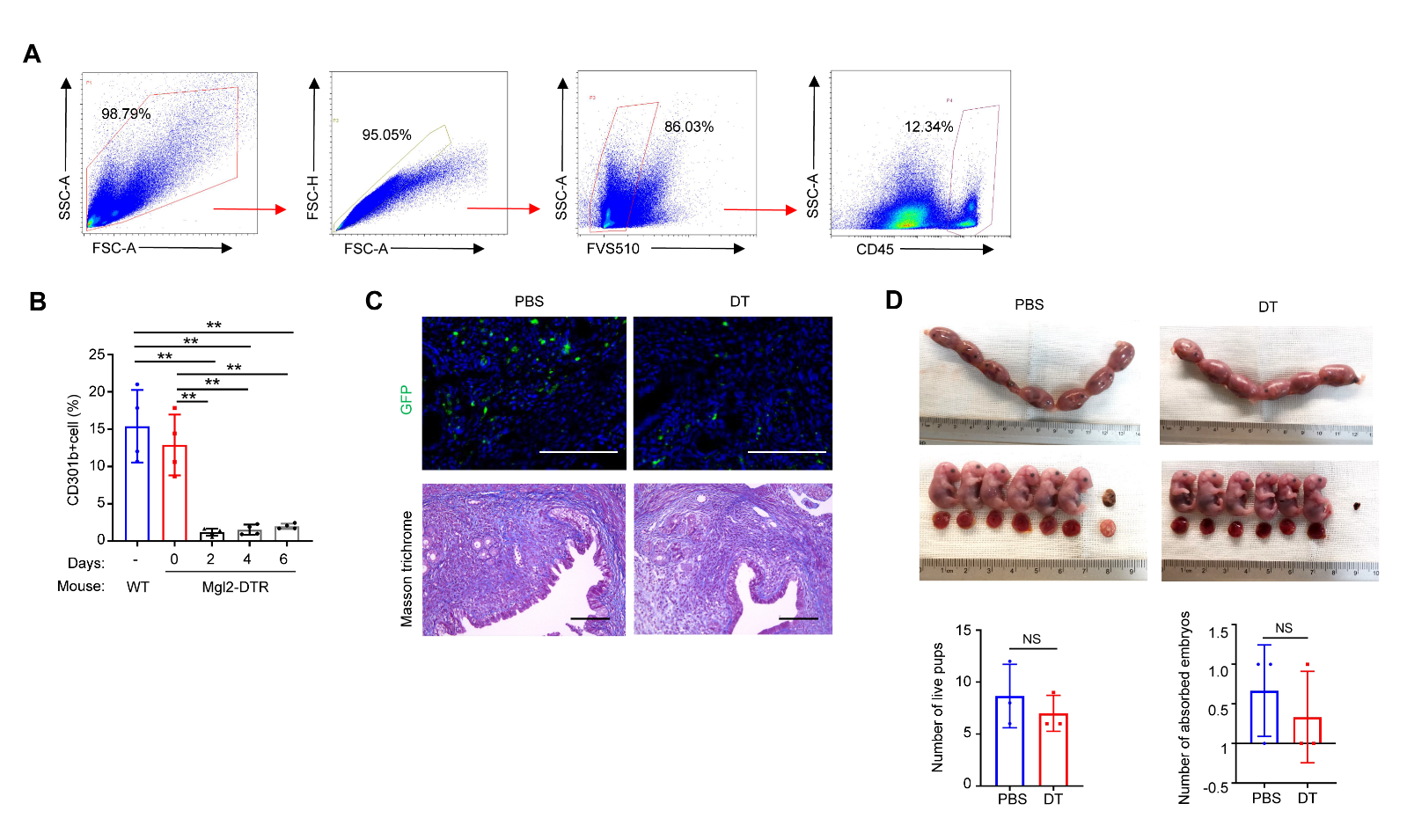
**Appendix Figure S7**

**Appendix Figure S7. Effects of depletion of CD301b+ macrophages on the fibrotic phenotype and the number of live pups and absorbed embryos in mice.**

A. Flow cytometry gating strategy for FVS510-CD45+ cells of murine uterine cells.
B. Flow cytometric analysis of proportion of CD301b+ cells for 0, 2, 4, 6 days after single injection of DT in the uterus of Mgl2-DTR and Wild-type mice (n=4 mice for each group).
C. Representative images of GFP-MGL2 expression and Masson trichrome staining in endometrium of MGL2-DTR mice injected by DT or PBS four days later (n=3 mice for each group). Scale bar, 100μm.
D. The number of live pups and absorbed embryos at day 18.5 of pregnancy (18.5 dpc) in mice of PBS and DT group (n=3 mice for each group).

Data are presented as mean ± SEM. (B) One-way ANOVA with Tukey’s post hoc analysis. (D) Two-tailed Student’s t-test. NS, not significant.

**Appendix Figure S8**

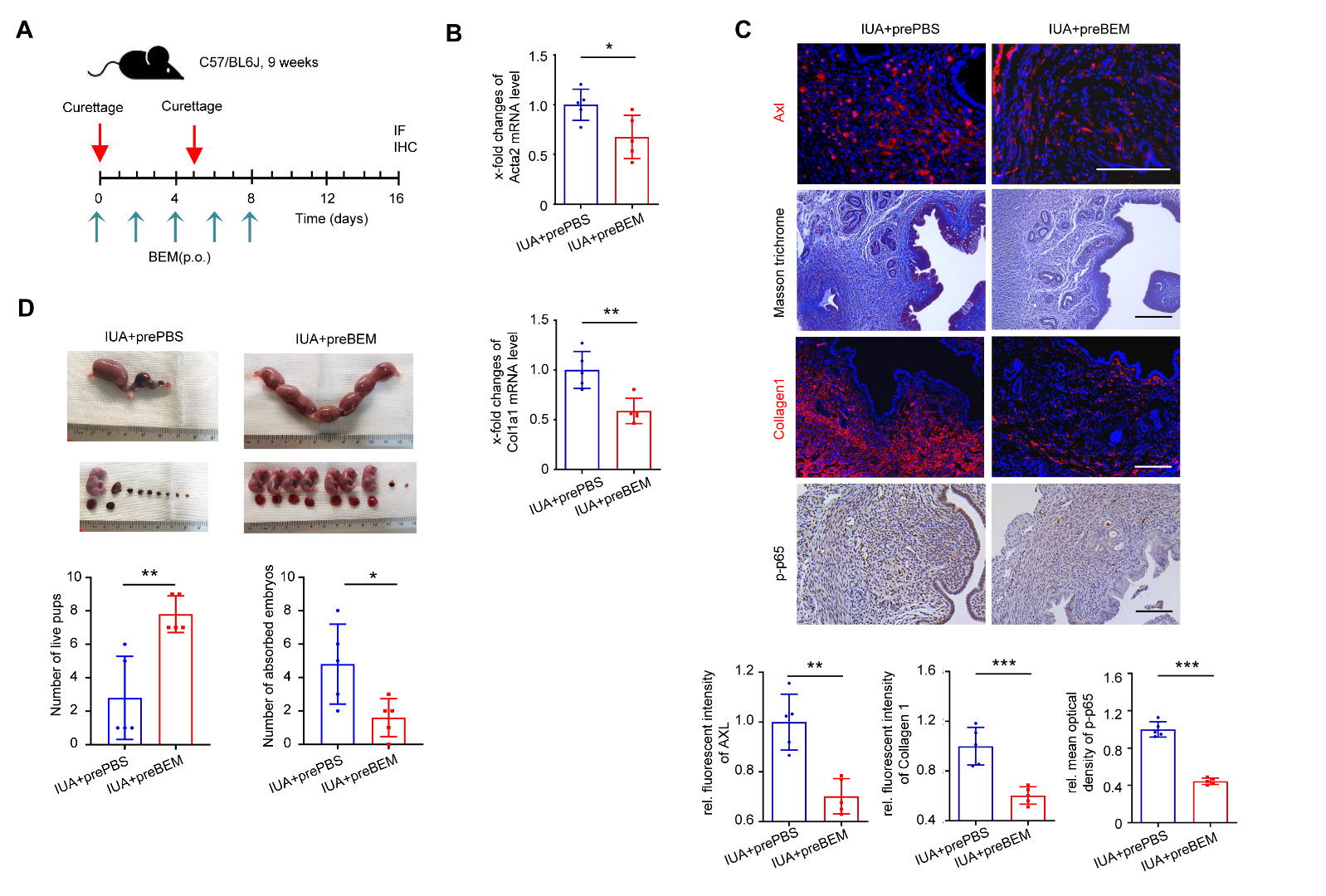

**Appendix Figure S8. Preventive (pre) effects of** [**Bemcentinib**](https://www.medchemexpress.cn/R428.html) **on endometrial fibrosis.**

A. Outline of the design of Bemcentinib (BEM) administration during the model induction.
B. qRT-PCR analysis of Acta2 and Col1a1 expression in the uterus of IUA+prePBS and IUA+preBEM mice (n=5 mice for each group).
C. Representative images of immunostaining for AXL, Collagen1, p-p65 and Masson trichrome staining in the endometrium of IUA+prePBS group and IUA+preBEM group (n=5 mice for each group).
D. The number of live pups and absorbed embryos at 18.5 dpc between IUA+prePBS group and IUA+preBEM group (n=5 mice for each group).

Scale bar: 100μm. Data are presented as mean ± SEM. (B-D) Two-tailed Student’s t-test. *, P<0.05; **, P<0.01; ***, P<0.001.

**Appendix Table S1** **- Clinical characteristics of women included in scRNA-seq analysis ^a)^.**

|  | Control-1 | Control-2 | Control-3 | Patient-1 | Patient-2 | Patient-3 |
| --- | --- | --- | --- | --- | --- | --- |
| Age (year) | 29 | 31 | 27 | 25 | 33 | 28 |
| Times for D&C | 0 | 0 | 0 | 2 | 5 | 3 |
| Times for TCRA | 0 | 0 | 0 | 3 | 3 | 1 |
| Parity (time) | 0 | 0 | 0 | 0 | 0 | 0 |
| Length of menstrual cycle (day) | 28 | 30 | 30 | 31 | 30 | 30 |
| Day after menstruation (day) | 13 | 13 | 14 | 14 | 14 | 14 |
| Follicle size on the day of hysteroscopy (mm) | 18*18*17 | 17*16*17 | 16*16*18 | 16*17*16 | 17*18*17 | 16*18*7 |
| EM on the day of hysteroscopy (mm) | 8.4 | 8.1 | 9.7 | 5.8 | 6.4 | 6.1 |
| Serum progesterone (nmol/L) | 0.34 | 0.14 | 0.21 | 0.21 | 0.31 | 0.35 |
| AFS score | 0 | 0 | 0 | 12 | 10 | 10 |

a) D&C, dilation and curettage; TCRA, transcervical resection of adhesion; EM, endometrial thickness; AFS: American Fertility Society.

**Appendix Table S2** **- Cell numbers of each type of cells in scRNA-seq analysis.**

| Cluster | Control-1 | Control-2 | Control-3 | Patient-1 | Patient-2 | Patient-3 |
| --- | --- | --- | --- | --- | --- | --- |
| Stromal cell | 5006 | 5911 | 7621 | 6984 | 4587 | 5766 |
| Epithelial cell | 1288 | 1498 | 1466 | 1208 | 935 | 1441 |
| Myofibroblast-like cell | 434 | 517 | 527 | 750 | 466 | 592 |
| Endothelial cell | 302 | 216 | 237 | 412 | 299 | 289 |
| T cell | 302 | 927 | 1127 | 1109 | 510 | 1343 |
| Macrophages/DC | 247 | 339 | 311 | 243 | 192 | 275 |
| Natural killer cell | 115 | 336 | 629 | 175 | 152 | 467 |
| B cell | 19 | 20 | 21 | 10 | 10 | 12 |
| Mast cell | 12 | 26 | 24 | 10 | 33 | 22 |
| Total | 7725 | 9790 | 11963 | 10901 | 7184 | 10207 |

**Appendix Table S3 - Cell numbers of macrophage/DC in scRNA-seq analysis.**

| Cluster | Control-1 | Control-2 | Control-3 | Patient-1 | Patient-2 | Patient-3 |
| --- | --- | --- | --- | --- | --- | --- |
| Macrophage | 233 | 302 | 281 | 227 | 169 | 251 |
| DC | 14 | 37 | 30 | 16 | 23 | 24 |

**Appendix Table S4** **- Patient clinical information ^b)^.**

| Items | Control | Patient | *P* value |
| --- | --- | --- | --- |
|  | (n = 48) | (n = 48) |  |
| Age (year) | 30.29 ±0.60 | 31.04 ± 0.77 | > 0.05 |
| Times for D&C | 1.50 ± 0.24 | 2.96 ± 0.23 | < 0.05 |
| Times for TCRA | 0 | 2.18 ± 0.21 | - |
| Maximum endometrial thickness (mm) | 10.23±0.38 | 6.49±0.33 | < 0.05 |
| Length of menstrual cycle (day) | 30.21±0.23 | 30.59±0.53 | > 0.05 |
| Day after menstruation | 12.18±0.41 | 13.04±0.41 | > 0.05 |
| Serum progesterone(day) | 0.43±0.12 | 0.41±0.59 | > 0.05 |
| AFS score | 0 | 10.21±0.26 | < 0.05 |
| Diagnosis | Tubal infertility | Intrauterine adhesion |  |

b) D&C, dilation and curettage; TCRA, transcervical resection of adhesion; AFS, American Fertility Society.

**Appendix Table S5 -** **Primer sequences.**

| Gene name | Forward Primer (5'-3') | Reverse Primer (5'- 3') |
| --- | --- | --- |
| *18S rRNA* | CTTTGGTCGCTCGCTCCTC | CTGACCGGGTTGGTTTTGAT |
| *AXL* | GTGGGCAACCCAGGGAATATC | GTACTGTCCCGTGTCGGAAAG |
| *COL1A1* | GAGGGCCAAGACGAAGACATC | CAGATCACGTCATCGCACAAC |
| *TGFB1* | CAAGCAGAGTACACACAGCAT | TGCTCCACTTTTAACTTGAGCC |
| *TIMP1* | ACCACCTTATACCAGCGTTATGA | GGTGTAGACGAACCGGATGTC |
| *NFKB1* | GAAGCACGAATGACAGAGGC | GCTTGGCGGATTAGCTCTTTT |
| *CXCL8* | ACTGAGAGTGATTGAGAGTGGAC | AACCCTCTGCACCCAGTTTTC |
| *IL1B* | ATGATGGCTTATTACAGTGGCAA | GTCGGAGATTCGTAGCTGGA |
| *IL18* | TCTTCATTGACCAAGGAAATCGG | TCCGGGGTGCATTATCTCTAC |
| *ACTA2* | CTATGAGGGCTATGCCTTGCC | GCTCAGCAGTAGTAACGAAGGA |
| *Col1a1* | CTGGCGGTTCAGGTCCAAT | TTCCAGGCAATCCACGAGC |
| *Acta2* | CCCAGACATCAGGGAGTAATGG | TCTATCGGATACTTCAGCGTCA |

**Appendix Table S6 - Antibodies used for immunohistochemistry (IHC), immunofluorescence (IF), western blot (WB), Flow cytometry (FC).**

| **Antibodies** | **SOURCE** | **IDENTIFIER** |
| --- | --- | --- |
| Anti-mouse CD45 APC-Cy7 | BD Biosciences | Cat# 557659, clone 30-F11; RRID: AB_396774 |
| Anti-mouse CD11b FITC | BD Biosciences | Cat#557396, clone M1/70; RRID: AB_396679 |
| Anti-mouse F4/80 Alexa Fluor 647 | BD Biosciences | Cat#565853, clone T45-2342; RRID: AB_2744474 |
| Anti-mouse CD301b PE | Biolegend | Cat#146803, clone URA-1; RRID: AB_2562943 |
| Rat IgG2a, κ PE | Biolegend | Cat#400507, clone RTK2758 |
| FVS 510 | BD Biosciences | Cat#564406; RRID: AB_2869572 |
| Anti-human CD45 PE-Cy7 | BioLegend | Cat#368532, clone 2D1; RRID: AB_2715892 |
| Anti-human CD14 FTIC | BioLegend | Cat#301804, clone M5E2; RRID: AB_314186 |
| Anti-human CD80 PE | BioLegend | Cat#305208, clone 2D10; RRID: AB_314504 |
| Mouse IgG1, κ PE | BioLegend | Cat#400111, clone MOPC-21 |
| Anti-human CD163 BV650 | BD Biosciences | Cat#563888, clone GHI/61; RRID: AB_2738468 |
| Mouse IgG1, κ BV650 | BD Biosciences | Cat#563231, clone X40; RRID: AB_2869470 |
| Anti-human CD301 APC | BioLegend | Cat#354706, clone H039G3; RRID: AB_11219389 |
| Mouse IgG2a, κ APC | BioLegend | Cat#400221, clone MOPC-173; RRID: AB_2891178 |
| Goat anti-GAS6 | R&D Systems | Cat# AF885; RRID: AB_2108079 |
| Goat anti-GAS6 | R&D Systems | Cat# AB885; RRID: AB_354376 |
| Goat anti-GAS6 | R&D Systems | Cat# AF986; RRID: AB_2263130 |
| Rabbit anti-CCL8 | Abcam | Cat# ab9671; RRID: AB_308749 |
| Rabbit anti-AXL | Cell Signaling Technology | Cat#8661, clone C89E7; RRID: AB_11217435 |
| Rabbit anti-Phospho-AXL (Tyr779) | Cell Signaling Technology | Cat#96453 |
| Mouse anti-AXL | Abcam | Cat# ab89224, clone MM0098-2N33; RRID: AB_2049189 |
| Goat anti-AXL | R&D Systems | Cat# AF854; RRID: AB_355663 |
| Mouse anti-CD14 | Abcam | Cat# ab182032, clone 4B4F12 |
| HRP-rabbit anti-β-actin | Abclonal | Cat# AC028; RRID: AB_2769861 |
| Rabbit anti-MERTK | Abcam | Cat# ab52968, clone Y323; RRID: AB_2143584 |
| Rabbit anti-Phospho-MERTK (Y749+Y753+Y754) | Abcam | Cat#ab14921; RRID: AB_2250636 |
| Rabbit anti-TYRO3 | Abcam | Cat# ab109231, clone EPR4308 |
| Rabbit anti-Phospho-TYRO3 (Tyr681) | Biorbyt | Cat#orb186274 |
| Mouse anti-CD68 | Abcam | Cat# ab955, clone KP1; RRID: AB_307338 |
| Rabbit anti-CD301 | Novus | Cat# NBP1-84591; RRID: AB_11031743 |
| Mouse anti-α-SMA | Abcam | Cat# ab7817, clone 1A4; RRID: AB_262054 |
| Rabbit anti- α-SMA | Abcam | Cat# ab5694; RRID: AB_2223021 |
| Mouse anti-Collagen 1 | Abcam | Cat# ab6308, clone COL-1; RRID: AB_305411 |
| Rabbit anti-Collagen 1 | Proteintech | Cat# 14695-1-AP; RRID: AB_2082037 |
| Rabbit anti-CD10 | Abcam | Cat# ab256494, clone EPR22867-118; RRID: AB_2894853 |
| Rabbit anti-Phospho-NF-κB p65 (Ser 536) | Cell Signaling Technology | Cat#3033, clone 93H1; RRID: AB_331284 |
| Rabbit anti- NF-κB p65 | Cell Signaling Technology | Cat#8242, clone D14E12; RRID: AB_10859369 |
| Rabbit anti-Phospho-NF-κB p65 (Ser 536) | Abcam | Cat#ab86299; RRID: AB_1925243 |
| Goat anti-CD301a/b | R&D Systems | Cat#AF4297; RRID: AB_2248147 |
| Rabbit anti-Fibronectin | Abcam | Cat#ab2413; RRID: AB_2262874 |
| Rabbit anti-CTGF | Cell Signaling Technology | Cat#86641; RRID: AB_2800085 |
| Rabbit anti-CD24 | Abcam | Cat#ab278509 |
| Rabbit anti-LUM | Abcam | Cat#ab168348; RRID: AB_2920864 |
| Alexa Fluor 594 Donkey anti-rabbit | Abcam | Cat#ab150076; RRID: AB_2782993 |
| Alexa Fluor 594 Donkey anti-mouse | Abcam | Cat#ab150108; RRID: AB_2732073 |
| Alexa Fluor 488 Donkey anti-rabbit | Abcam | Cat# ab150073; RRID: AB_2636877 |
| Alexa Fluor 488 Donkey anti-mouse | Abcam | Cat# ab150105; RRID: AB_2732856 |
| Alexa Fluor 488 Donkey anti-goat | Abcam | Cat# ab150129; RRID: AB_2687506 |
| Alexa Fluor 594 Donkey anti-goat | Abcam | Cat# ab150132; RRID: AB_2810222 |
| Alexa Fluor 488 Goat anti-rat | Abcam | Cat# ab150157; RRID: AB_2722511 |
